# Neural Dissociation of Cognitive Maps and Graphs in the Human Brain

**DOI:** 10.64898/2026.10.09.757922

**Authors:** Yangwen Xu, Max A. B. Hinrichs, Roberto Bottini, Christian F. Doeller

## Abstract

Cognitive maps and graphs offer two distinct ways of organizing knowledge: maps arrange items within a spatial layout, whereas graphs thread them together through experiential transitions. However, spatial proximity and experiential contiguity are typically correlated, and content represented in one format can often be transformed into the other, making their neural representations difficult to disentangle. Here, we orthogonally manipulated Euclidean distance, indexing spatial proximity, and shortest-path distance, indexing experiential contiguity, within the same learned environment. Using fMRI and whole-brain searchlight representational similarity analysis, we identified dissociable representations of these two formats in the human brain. Euclidean distance was represented in the intraparietal sulcus, whereas shortest-path distance was represented in the hippocampal/parahippocampal region and in cortical areas that overlapped substantially with scene-selective regions. ROI analyses further supported the specificity of these representations across regions. Together, these findings provide neural evidence for a dissociation between cognitive maps and cognitive graphs.

## Introduction

Cognitive maps and graphs are two fundamental forms of mental representation (Warren, 2019; Peer et al., 2021; Figure 1A). In a Euclidean map, items are arranged in a low-dimensional space; their identities are defined by their locations. In a cognitive graph, items are represented as nodes connected by edges; their identities are defined through their relations. These two forms of mental representation are not only useful for navigation but also serve as built-in mental structures for organizing knowledge across a wide range of cognitive domains (Behrens et al., 2018; Bellmund et al., 2018; Peer et al., 2021).

**Figure 1.**
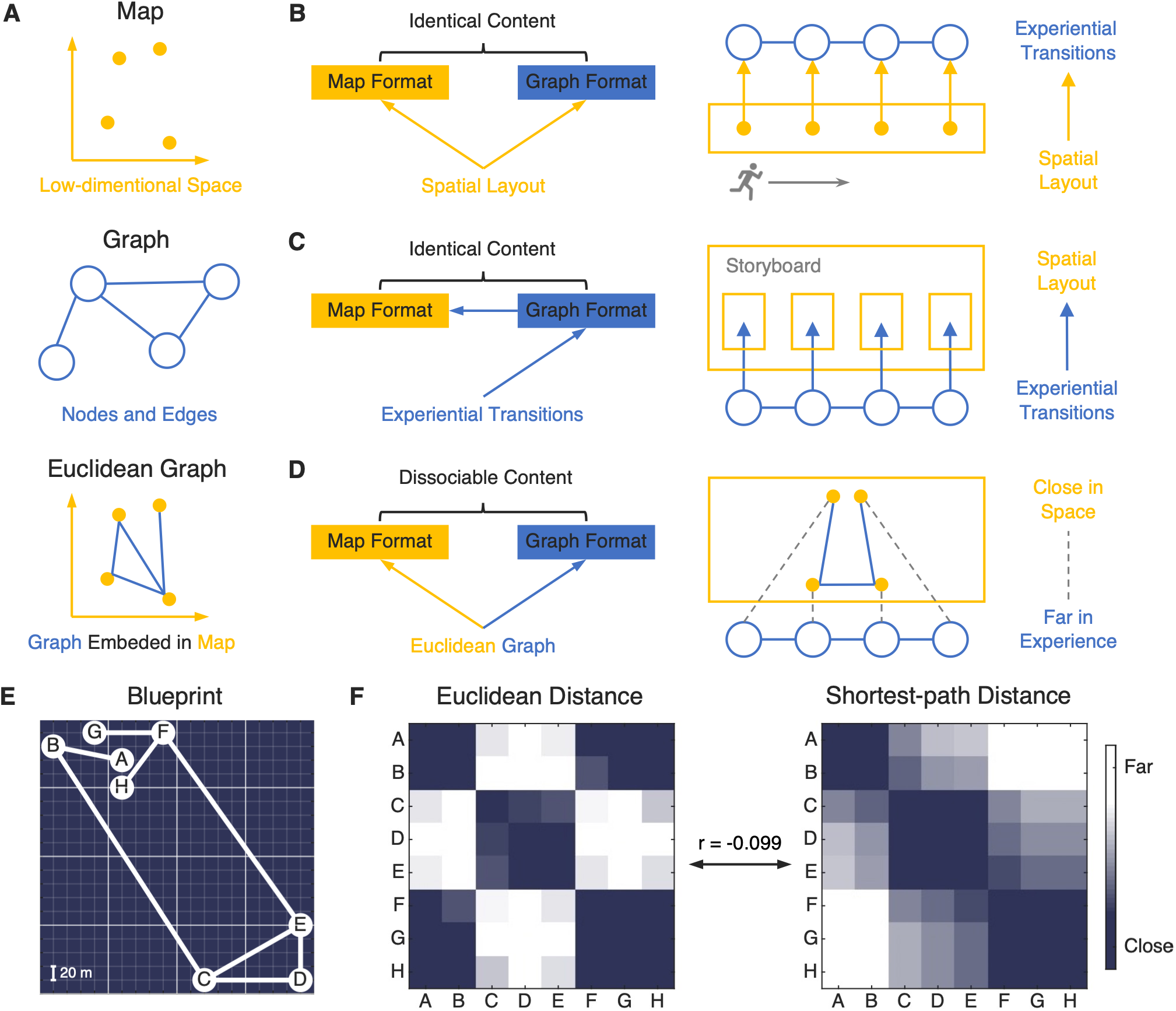
Experimental design. **(A)** Two forms of mental representation. A map arranges items in low-dimensional Euclidean space, and a graph organizes items according to their relationships. A Euclidean graph is a graph embedded in Euclidean space. **(B)** Spatial layouts can be “relationalised” in a graph format: during navigation, locations are encoded as nodes and movements between them as edges, recasting the spatial layout into experiential transitions. **(C)** Experiential transitions can be “spatialized” in a map format: when illustrating a story plot, time-adjacent events are arranged in close spatial proximity (e.g., a storyboard), translating experiential transitions into a spatial layout. **(D)** Map and graph representations can be dissociable in a Euclidean graph: close places can be far away in experience (i.e., “so close yet so far”). To maintain information with different content, neural systems prioritize information in their own formats and limit cross-format transfer. Thus, we can disentangle these two types of representations based on their content. **(E)** Blueprint of the virtual environment: a 400 × 400 m plane comprising eight locations (A–H) connected by routes. **(F)** Representational dissimilarity matrix of the virtual environment: map distance (Euclidean) and graph distance (shortest-path) were manipulated to be orthogonal across the 28 pairs (Pearson’s r = −0.099).

It remains debated what the neural underpinnings of these two forms of mental representation are and whether they are dissociable (Peer et al., 2021). As an illustrative example, the hippocampus has been a central focus of this debate (Eichenbaum et al., 1999; Lisman et al., 2017). In their seminal book, O’Keefe and Nadel (1978) proposed that the hippocampus represents a cognitive map corresponding to an absolute, unified space. This proposal was motivated by the discovery of hippocampal place cells, which fire when an animal occupies a specific location within an environment (O’Keefe & Dostrovsky, 1971). In contrast, Eichenbaum and Cohen (2001) proposed the Relational Memory Theory, which posits that the hippocampus represents the relations among the constituent elements of experience—a view consistent with a cognitive graph. Eichenbaum (2014) argued that the same hippocampal place cells can also function as time cells, with their sequential activation reflecting transitions in experience rather than absolute spatial locations (see related sequence- and transition-based accounts by Stachenfeld et al., 2017; Buzsáki & Tingley, 2018; George et al., 2021).

The crux of this issue lies in the fact that the same content can be encoded by both forms of mental representation, making it difficult to attribute observed neural activity uniquely to one or the other. For example, spatial layouts can be “relationalised” in a graph format, as proximal locations become related through experience when an agent traverses space (Figure 1B). Conversely, experiential transitions can be “spatialized” in a map format, as related experiences become spatially proximal when elements of experience are mentally arranged within a spatial layout (Figure 1C; on the spatial construal of time, see Lakoff & Johnson, 1980; Boroditsky, 2000).

One approach to address this challenge is to have the brain learn a Euclidean graph that jointly specifies node locations and edge connections (Figure 1A). Within this framework, spatial layout and experiential transitions are both present and can be manipulated orthogonally to one another. For instance, two locations may be spatially proximal yet distant in experience due to the absence of a direct connecting path. Because orthogonalized map and graph information are both available within the same environment, the corresponding neural systems should be preferentially engaged in representing information in their respective formats, thereby blocking cross-format transfer and minimizing confounding (Figure 1D).

This fMRI study applies this approach to disentangle neural representations of cognitive maps and graphs in the human brain. We designed a virtual environment consisting of eight locations (nodes) connected by paths (edges), with each location marked by a unique monument (Figure 1E; Figures S1 and S2). Euclidean distance was used to quantify distance in the cognitive map, whereas shortest-path distance was used to quantify distance in the cognitive graph. These two distance measures were orthogonally manipulated across the 28 possible pairs of locations (Figure 1F; r = −0.099). Participants learned both map and graph knowledge of the environment concurrently from a first-person perspective (Figure 2A). They could visually perceive the Euclidean distances between monuments but were constrained to navigate only along the paths. This “open maze” environment ensured that participants could readily learn the spatial layout specified by the cognitive map, while their experiential transitions were governed by the structure of the cognitive graph (Peer et al., 2024). Using whole-brain searchlight representational similarity analysis (RSA), we asked, for each brain region, whether its activity patterns reflected map representations indexed by Euclidean distance or graph representations indexed by shortest-path distance (Kriegeskorte et al., 2006; Kriegeskorte et al., 2008).

**Figure 2.**
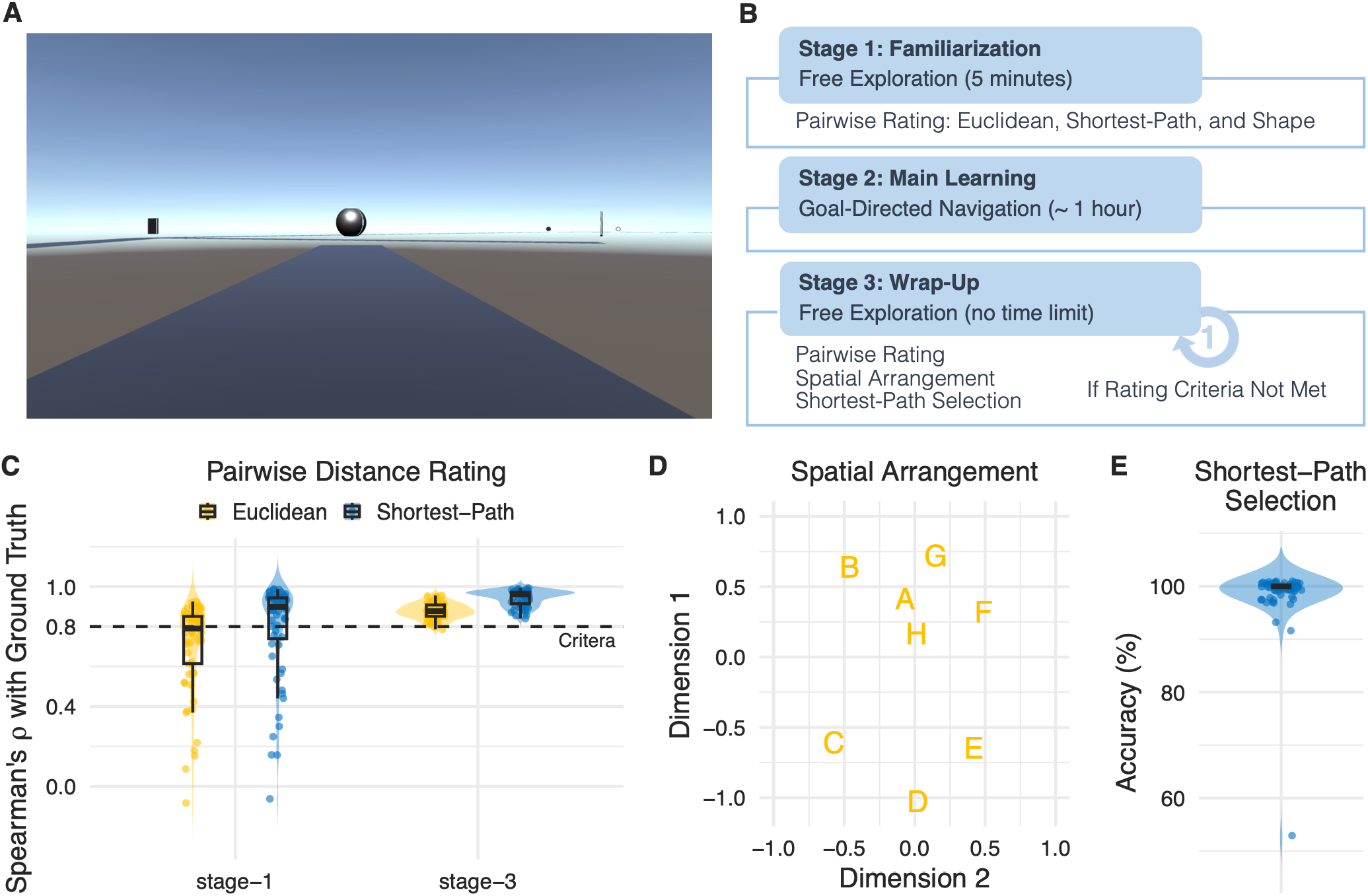
Virtual environment and behavioural results. **(A)** The virtual environment was an “open maze” presented from a first-person, ground-level perspective. It was “open” in that no visible obstacles blocked the view, allowing participants to perceive Euclidean distances. It remained a “maze” because invisible barriers prevented off-route traversal, restricting movement to predefined paths. **(B)** The learning procedure on the day before fMRI scanning comprised three stages, with behavioural tests assessing cognitive map and graph knowledge administered after Stages 1 and 3. If predefined performance criteria were not met after Stage 3, participants completed an additional free-exploration stage (see Methods for details). **(C)** Pairwise rating results: Spearman correlations between pairwise ratings and the ground truth across the 28 pairs after Stages 1 and 3. **(D)** Spatial arrangement task results: multidimensional scaling of the group-averaged pairwise distances from the spatial arrangement task. **(E)** Shortest-path selection task results: accuracy in the shortest-path selection task.

## Results

### Participants acquired both map and graph knowledge of the virtual environment

On the day prior to MRI scanning, participants learned the virtual environment across three stages (see Methods). They first completed 5 min of free exploration (Stage 1), followed by goal-directed navigation training (Stage 2), and then an untimed free exploration period (Stage 3). Behavioural assessments administered after Stages 1 and 3 measured participants’ acquisition of both map and graph knowledge (Figure 2B; Table S1).

Figure 2C shows the Spearman correlations between ground-truth distances and participants’ pairwise distance ratings, computed separately for Euclidean and shortest-path distances. To ensure adequate acquisition of both map and graph knowledge, participants repeated Stage 3 if either correlation was below 0.8 or if the correlation between Euclidean and shortest-path ratings exceeded 0.25. Overall, 77% of participants (47/61) met these criteria after the first Stage 3 assessment, and an additional 15% (9/61) met the criteria after repeating Stage 3. The remaining five participants did not meet the learning criteria, but their performance was close to the 0.8 correlation threshold (Figure 2C); therefore, these participants were retained in the analyses.

Figure 2D shows the multidimensional scaling (MDS) results from the spatial arrangement task administered after Stage 3, in which participants arranged monument icons on a computer screen according to their relative positions in the virtual environment. MDS was applied to the participant-averaged pairwise Euclidean distance matrix. The reconstructed spatial layout closely matched the actual layout of the virtual environment (Figure 1E).

Figure 2E shows performance on the shortest-path selection task administered after Stage 3, in which participants identified which of two candidate monuments lay along the shortest path between a pair of locations. All but one participant performed nearly perfectly, with the remaining participant performing at chance level. Because this participant performed well on the pairwise rating task, we interpreted their chance-level performance as likely reflecting a misunderstanding of the task instructions rather than insufficient cognitive graph knowledge. We therefore retained this participant in the analyses.

Together, these results indicate that participants acquired both map and graph knowledge of the virtual environment by the end of Stage 3 (Figure 2C–E).

In addition to the assessments of cognitive map and graph knowledge, participants also completed shape dissimilarity ratings for each pair of monuments (Figure S3A). The participant-averaged shape dissimilarity matrix obtained after Stage 3 served as the shape dissimilarity matrix for subsequent representational similarity analyses (Figure S3B).

### Dissociable hippocampal networks were engaged during memory recall and oddball detection

The fMRI task was designed to prompt participants to recall the map and graph knowledge associated with the eight locations, cued by images of the corresponding monuments (Figure 3A). During standard trials (90% of trials), participants were instructed to recall each monument’s position within the environment, including both its spatial location and its connections to other monuments, and to press a button with their right index finger once this information came to mind. During oddball trials (10% of trials), participants pressed a different button with their right middle finger when presented with a novel monument that was not part of the learned environment.

**Figure 3.**
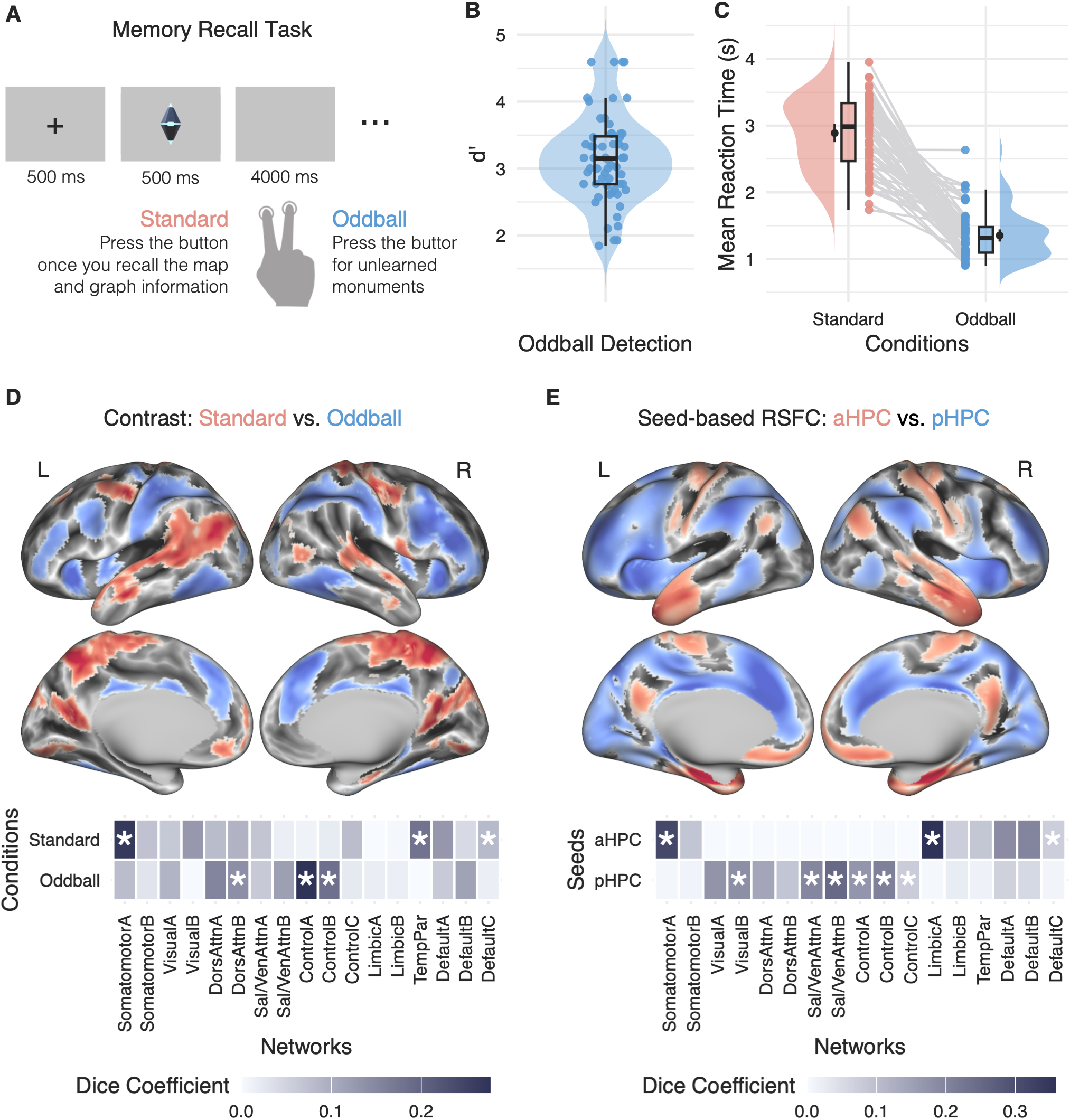
fMRI task design and hippocampal network engagement during memory recall and oddball detection. **(A)** Memory recall task and trial timing. **(B)** Oddball detection sensitivity, indexed by *d′*. **(C)** Mean reaction times for correct standard and oddball trials, computed within each participant. Points and connecting lines indicate individual participants; black points indicate group means, and error bars indicate 95% confidence intervals. **(D)** Task-activation contrasts and functional-network correspondence between the standard and oddball conditions. The z-statistic maps were thresholded using a voxel-wise cluster-forming threshold of *p* < 0.001 and a whole-brain cluster-level FWE-corrected threshold of *p* < 0.05 (L = left; R = right). The anterior and posterior hippocampi are highlighted in Figure S4. Network correspondence was assessed using the Yeo 17-network atlas (Yeo et al., 2011). Asterisks indicate statistically significant correspondence based on spin-test permutations. **(E)** Resting-state functional connectivity (RSFC) contrasts and functional-network correspondence between the anterior hippocampal (aHPC) and posterior hippocampal (pHPC) seeds. The seeds were defined from the significant hippocampal clusters identified in the standard > oddball and oddball > standard task-activation contrasts, respectively. The z-statistic maps representing the contrasts between the aHPC and pHPC RSFC maps were thresholded using whole-brain voxel-wise FWE correction at *p* < 0.05 (L = left; R = right). Network correspondence was assessed using the Yeo 17-network atlas (Yeo et al., 2011). Asterisks indicate statistically significant correspondence based on spin-test permutations.

The oddball trials served as an attention check. The high *d′* for oddball detection indicated that participants were attentive (M ± SD = 3.14 ± 0.65; Figure 3B). Reaction times were analysed using a linear mixed-effects model with trial type (standard vs. oddball) as a fixed effect and participant-specific random intercepts and slopes for trial type. This analysis showed that participants spent more time on standard trials than on oddball trials (F(1, 59.17) = 270.44, p < 0.001; Figure 3C), further suggesting that they engaged in recalling the map and graph information associated with the learned monuments.

Figure 3D shows the cortical distribution of the task-activation contrasts between the standard and oddball conditions, together with their correspondence with canonical functional brain networks (Yeo et al., 2011; Kong et al., 2025). Hippocampal effects are highlighted separately in Figure S4. The standard condition elicited significantly greater activation in the anterior hippocampus, the somatomotor cortex, the temporoparietal cortex, and regions of the default network. By contrast, the oddball condition elicited significantly greater activation in the posterior hippocampus and regions of attentional and control networks.

We also compared seed-based resting-state functional connectivity (RSFC) between anterior and posterior hippocampal seeds defined from the significant hippocampal clusters identified in the standard > oddball and oddball > standard contrasts, respectively (Figure 3E). The resulting connectivity differences showed a pattern broadly consistent with the task-activation contrasts. The anterior hippocampus exhibited stronger functional connectivity with the somatomotor cortex, limbic regions, and regions of the default network, whereas the posterior hippocampus exhibited stronger functional connectivity with regions of attentional and control networks.

These results are consistent with recent evidence for functional specialization along the longitudinal axis of the human hippocampus. The anterior hippocampus, which is preferentially connected with the default network, appears to be more strongly involved in memory recall, whereas the posterior hippocampus, which is preferentially connected with salience-related networks, appears to be more strongly involved in detecting salient or novel events (Angeli et al., 2025). This dissociation may help resolve the puzzle of why hippocampal activity follows a U-shaped pattern across a continuum from novelty through familiarity to recollection, with stronger responses to both novel and recollected information than to familiar information (Kim, 2021). The two arms of this U-shaped response may preferentially engage different regions along the hippocampal longitudinal axis, with the anterior hippocampus contributing more strongly to memory recall and the posterior hippocampus to novelty detection (Angeli et al., 2025; Quent et al., 2026).

### Dissociable brain regions represent cognitive maps and graphs

As a validity check, we first performed a whole-brain searchlight RSA using monument shape dissimilarity. This analysis revealed significant effects in brain regions associated with shape processing, including the bilateral inferolateral occipitotemporal cortex, bilateral intraparietal sulcus, and right ventral premotor cortex (Figure S5A). The spatial distribution of these effects closely corresponded to the shape-selective regions identified in our previous study using the univariate contrast of object shape > object function (Figure S5B; Xu et al., 2023).

Having established that the whole-brain searchlight RSA recovered the expected pattern of shape-related representations, we then applied the same approach to examine the neural representations of cognitive maps and graphs, indexed by Euclidean distance and shortest-path distance, respectively. The analyses revealed spatially dissociable patterns for the two representations (Figure 4A; Figure S6A). Euclidean distance was represented in the right intraparietal sulcus, whereas shortest-path distance was represented in three distinct regions: the right superior lateral occipital cortex, the right posteromedial region spanning the retrosplenial cortex and lingual gyrus, and the left hippocampal/parahippocampal region with an extension into the adjacent fusiform cortex.

**Figure 4.**
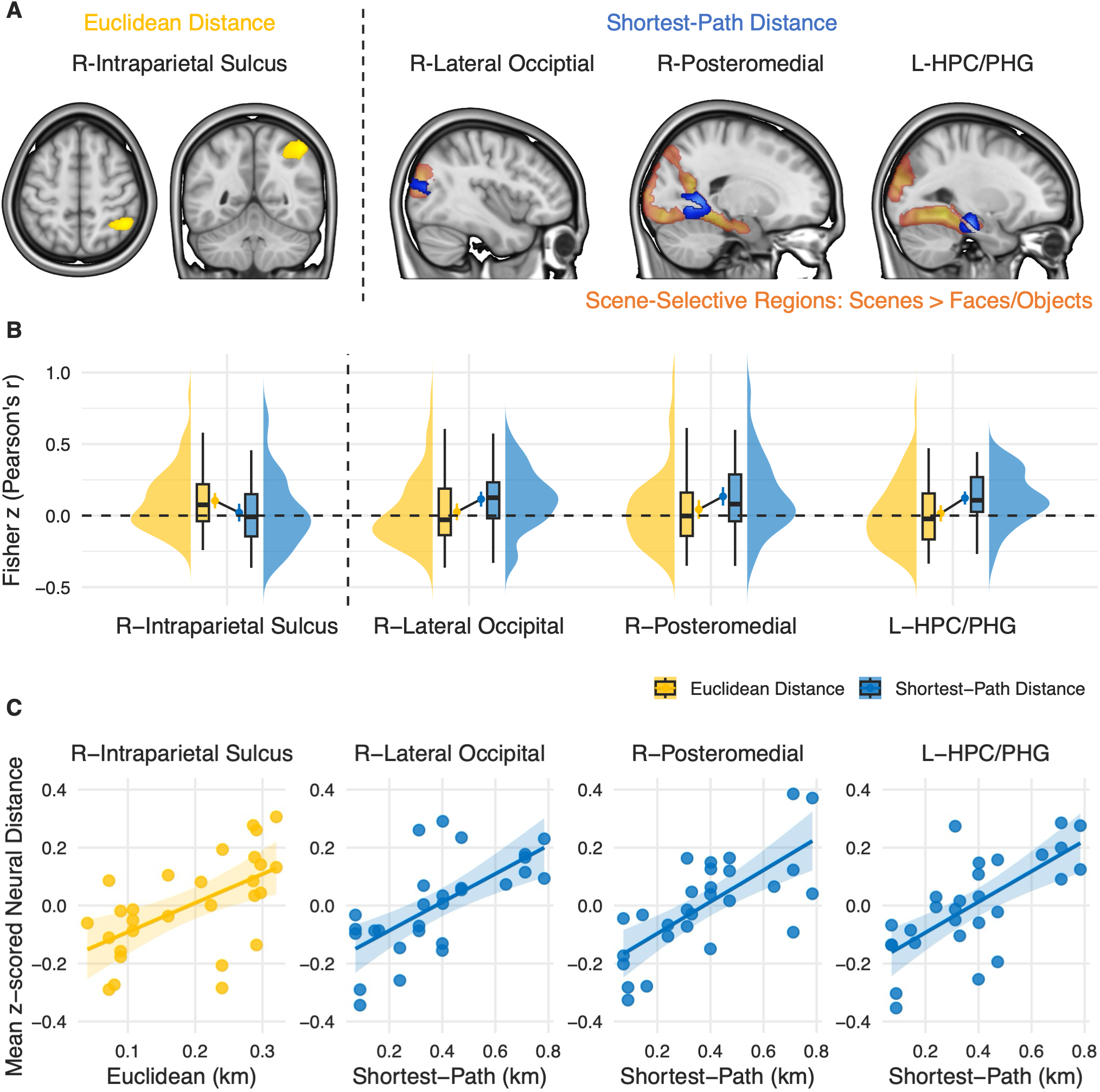
Neural representations of Euclidean and shortest-path distance. **(A)** Whole-brain searchlight RSA for Euclidean distance and shortest-path distance. Euclidean-distance representations were observed in the right intraparietal sulcus, whereas shortest-path-distance representations were observed in the right superior lateral occipital cortex, right posteromedial region, and left hippocampal/parahippocampal region. For the shortest-path-distance effects, scene-selective regions identified by the contrast scenes > faces/objects are shown in orange for anatomical comparison. Both searchlight RSA and scene-selective maps were thresholded at voxel-wise cluster-forming *p* < 0.001 and cluster-level FWE-corrected *p* < 0.05. **(B)** Representational profiles within regions identified in the whole-brain searchlight analysis. Participant-level Fisher *z*-transformed Pearson correlations are shown for Euclidean distance and shortest-path distance in the right intraparietal sulcus, right superior lateral occipital cortex, right posteromedial region, and left hippocampal/parahippocampal region. Points indicate group means and error bars indicate 95% confidence intervals. **(C)** Visualization of the relationships between model distance and neural distance within the searchlight-defined regions. Euclidean distance is shown for the right intraparietal sulcus, whereas shortest-path distance is shown for the right superior lateral occipital cortex, right posteromedial region, and left hippocampal/parahippocampal region. Neural distances were *z*-standardized across the 28 location pairs within each participant and then averaged across participants. Each point represents one of the 28 pairwise relationships among the eight locations. Lines show linear fits and shaded areas indicate 95% confidence intervals. Model distances are expressed in kilometres.

To characterize the representational profiles of these searchlight-defined regions, we next performed ROI-based RSA (Figure 4B). The right intraparietal sulcus showed a representation of Euclidean distance but no evidence for shortest-path distance (*t*(59) = 0.66, *p* = 0.51), whereas the three regions identified by the shortest-path searchlight showed the expected representation of shortest-path distance but no evidence for Euclidean distance (all uncorrected *p*s > 0.20). Consistent with this pattern, a 4 × 2 repeated-measures ANOVA with Region and Representation as within-participant factors revealed a significant Region × Representation interaction (*F*(3, 177) = 4.49, *p* = 0.005, *η_p_*^2^ = 0.071), indicating that the representations of Euclidean and shortest-path distance differed across regions. Because these ROIs were defined from the searchlight analysis itself, these analyses were used to characterize the representational profiles of the identified regions rather than as an independent test of the searchlight effects. To visualize these representational relationships, we additionally plotted neural distance against Euclidean distance in the right intraparietal sulcus and against shortest-path distance in the right superior lateral occipital cortex, right posteromedial region, and left hippocampal/parahippocampal region (Figure 4C). Across the 28 location pairs, greater model distance was descriptively associated with greater neural distance in the corresponding regions.

Despite counterbalancing the location–monument assignments across participants using a Latin-square design, rated pairwise shape dissimilarity still showed slight positive correlations with both Euclidean distance and shortest-path distance at the group level (Figure S3C). We therefore conducted a post-hoc robustness analysis to examine whether the observed effects could be explained by shared variance with monument shape. Partial correlations were computed for each variable while controlling for the other two variables (Figure S5C). The principal representational effects remained significant after Holm correction for multiple comparisons (all Holm-adjusted *p*s < 0.01), indicating that the observed Euclidean- and shortest-path-related representations were not readily attributable to their shared variance with shape dissimilarity.

Notably, several shortest-path effects were located in regions associated with scene processing (Figure 4A; Figure S6A). In particular, large proportions of the right superior lateral occipital cortex and the right posteromedial region fell within scene-selective regions defined by the contrast of scenes > faces/objects, with 81.03% and 68.41% of their voxels falling within these regions, respectively. We therefore further examined independently defined, functionally localized scene-selective regions (Figure S6B). ROI analyses were conducted separately in each hemisphere for the parahippocampal place area, retrosplenial cortex, and occipital place area, testing representations of Euclidean distance, shortest-path distance, and shape dissimilarity within each region (Figure S6C). Shortest-path distance was significantly represented in the right parahippocampal place area after correction for multiple comparisons, whereas the remaining effects did not survive correction. These findings suggest that graph-related information is represented within parts of the scene-selective system, although the spatial distribution of shortest-path effects does not correspond exactly to the boundaries of conventionally defined scene-selective regions.

Finally, to zoom in on the hippocampal–entorhinal system, we performed a searchlight RSA within a cytoarchitectonically defined search volume encompassing the hippocampal formation and entorhinal cortex (Figure S7). Within this restricted search volume, only shortest-path distance showed a significant effect, confined to the left hippocampal formation; no significant effects were observed for Euclidean distance or shape dissimilarity (Figure S7A). This pattern was consistent with the whole-brain searchlight analysis, which likewise identified a shortest-path-distance effect in the left hippocampal/parahippocampal region. We then examined the entorhinal cortex more specifically using cytoarchitectonically defined left and right entorhinal ROIs. These analyses revealed no significant representation of Euclidean distance, shortest-path distance, or shape dissimilarity in either hemisphere (all uncorrected *p*s > 0.25; Figure S7B).

## Discussion

In this study, we disentangled the neural representations of cognitive maps and graphs by orthogonally manipulating Euclidean distance and shortest-path distance within the same learned environment. Participants acquired both map and graph knowledge of the environment before scanning and were instructed to recall both forms of knowledge during scanning. Whole-brain searchlight RSA revealed spatially dissociable representations of the two formats: Euclidean distance was represented in the right intraparietal sulcus, whereas shortest-path distance was represented in the right superior lateral occipital cortex, right posteromedial region, and left hippocampal/parahippocampal region. These regional differences were further supported by distinct representational profiles across the identified regions, and the principal effects remained significant after accounting for shared variance with monument shape.

The hippocampal finding is particularly relevant to the longstanding debate over the nature of hippocampal representation. Since the discovery of place cells (O’Keefe & Dostrovsky, 1971) and the landmark case of patient H.M. (Scoville & Milner, 1957), there has been little doubt that the hippocampus represents both spatial and episodic content. The central question therefore concerns its underlying representational format—whether it uses space to arrange episodes in memory (i.e., a cognitive map) or uses memory to bind locations in space (i.e., a cognitive graph; Eichenbaum et al., 1999). Previous studies have reported that hippocampal representations reflect both map and graph structures (e.g., Morgan et al., 2011; Howard et al., 2014; Schapiro et al., 2016; Garvert et al., 2017; Ottink et al., 2025). However, in most studies, either only one form of information is available, allowing the other to be inferred from it, or both forms are available but correlated. In either case, the two representational formats are difficult to distinguish. Here, we orthogonally manipulated Euclidean and shortest-path distance within the same environment, making both forms of information simultaneously available. Under these conditions, the hippocampal/parahippocampal region represented shortest-path distance but showed no evidence for Euclidean distance, supporting a graph-like relational organization of hippocampal representations. Consistent with this perspective, in rodents, hippocampal neurons can distinguish different memory episodes occurring at the same physical location but on different trajectories (e.g., Wood et al., 2000). In humans, hippocampal damage can leave short-term path integration intact (Shrager et al., 2008), while disrupting the temporal organization of episodic memories (Dede et al., 2016). Moreover, the hippocampus is well suited for the neural implementation of a cognitive graph, with ensembles as nodes and modifiable synaptic connections between them constituting edges (Muller et al., 1996). Rapid forms of hippocampal synaptic plasticity, including behavioural-timescale synaptic plasticity, provide a plausible mechanism for establishing and updating these edges through experience (Bittner et al., 2017; Li et al., 2024).

Beyond the hippocampus, graph-like representations also emerged in scene-selective regions. The shortest-path effects in the right superior lateral occipital and posteromedial regions substantially overlapped with scene-selective activations. The superior lateral occipital effect overlapped with the occipital place area (OPA; Dilks et al., 2013), whereas the posteromedial effect overlapped with the parahippocampal place area (PPA; Epstein & Kanwisher, 1998) and retrosplenial cortex (RSC; Epstein et al., 2007). As the whole-brain effects did not perfectly coincide with the canonical boundaries of these scene-selective regions, follow-up ROI analyses revealed a significant shortest-path representation only in the right PPA. While scene-selective regions were traditionally characterized primarily in terms of local scene representation—the nodes (see Epstein & Baker, 2019, for review), accumulating evidence suggests that they also represent connections between local scenes along navigable routes—the edges (see Peer et al., 2021, for review). In the OPA, multivoxel activity in human fMRI represents navigational affordances for movement through the local scene (Bonner & Epstein, 2017). In the RSC, neural activity in rodents reflects an animal’s position within a route and the location of that route within the broader environment, while also encoding route distance and route structure (Alexander & Nitz, 2015, 2017; see Alexander et al., 2023, for review). As for the PPA, previous accounts have mostly emphasized its role in representing the spatial layout of the local scene (Epstein & Kanwisher, 1998). However, because local scenes are continuously linked by adjacent paths as we traverse an environment, our shortest-path effect suggests that the PPA may encode not only the spatial layout of individual scenes but also their experiential contiguity during navigation. Given the strong connectivity between the hippocampus and parts of the scene-selective system such as the RSC (e.g., Baldassano et al., 2016; Tullo et al., 2023), these two systems may work together to capture experiential transitions from an internal route perspective (Siegel & White, 1975), with the hippocampus indexing scene representations in scene-selective cortex.

By contrast, we found that the right IPS represented Euclidean distance but showed no evidence for shortest-path distance, suggesting that it supports a map-like representation. Unlike the hippocampus, the IPS is well suited for the neural implementation of a cognitive map. Its cortical organization is itself map-like, with locations in visual space systematically represented across multiple retinotopic maps on the cortical surface (Sereno et al., 2001; Swisher et al., 2007; Silver & Kastner, 2009). These retinotopic maps form part of the dorsal visual pathway, which specializes in spatial processing for visually guided action (Mishkin et al., 1983; Goodale & Milner, 1992). Neural recordings in the monkey lateral intraparietal area (LIP) have shown that multiple locations in space are represented simultaneously, with stronger responses to locations that are currently important for the animal’s behaviour, supporting the idea that LIP contains a priority map of space (Gottlieb et al., 1998; see Bisley & Goldberg, 2010, for review). fMRI studies in the human IPS further indicate that parietal maps can be engaged through top-down attention and the internal maintenance of spatial information, rather than being driven solely by bottom-up visual input (Corbetta et al., 2002; Silver et al., 2005; see Corbetta & Shulman, 2002, for review). Neuropsychological evidence suggests that bilateral occipitoparietal lesions can produce Bálint’s syndrome, characterized by simultanagnosia, oculomotor apraxia, and optic ataxia, leaving patients unable to perceive multiple elements as a coherent spatial whole or efficiently orient and act within the space in front of them (Bálint, 1909; Rizzo & Vecera, 2002; Chechlacz & Humphreys, 2014). Together, these findings suggest that parietal maps may provide a neural substrate for representing an environment from an external survey perspective (Siegel & White, 1975)—as a literal map laid out before the mind’s eye, in which multiple locations are maintained simultaneously, and task-relevant locations are prioritized. In our task, recalling a monument may therefore reinstate its corresponding location within this spatial representation. Monuments that are close in Euclidean space would therefore recruit more similar spatial codes in the IPS, providing a potential neural basis for the Euclidean-distance effect observed here. Because the visuospatial input was the monument image, the additional representation of monument shape in the IPS is also consistent with its role in representing visuospatial information from an external perspective.

Does the brain also maintain a map-like representation from an internal perspective within the environment? Such a representation would be distinct from both a graph representation from an internal route perspective associated with the hippocampus and a map representation from an external survey perspective associated with the IPS. The entorhinal cortex is a natural candidate for such a representation. While hippocampal place cells remap across environments, entorhinal grid cells exhibit periodic firing fields arranged in hexagonal lattices that shift or rotate coherently across environments (Hafting et al., 2005; Fyhn et al., 2007). One influential account proposes that grid-cell representations arise from continuous-attractor dynamics, in which recurrent connectivity constrains population activity to a low-dimensional manifold that serves as a continuous spatial map, while the population state shifts smoothly across this manifold as an individual moves through the environment (McNaughton et al., 2006; Burak & Fiete, 2009). Consistent with this account, population recordings have provided evidence for low-dimensional continuous-attractor dynamics (Yoon et al., 2013). However, neither the zoom-in searchlight nor the ROI analyses revealed evidence for Euclidean distance in the entorhinal cortex. Several possibilities may account for this null effect. First, entorhinal spatial representations may not be well captured by a model assuming that neural dissimilarity increases monotonically with physical Euclidean distance. Population activity within individual grid-cell modules has been shown to lie on a toroidal manifold (Gardner et al., 2022). Therefore, locations that are widely separated in physical space can be represented by similar population states, potentially obscuring a simple Euclidean-distance effect. Second, although grid-like representations have been reported in humans (Doeller et al., 2010; Jacobs et al., 2013), much evidence for continuous-attractor dynamics comes from rodents. Given the prominent role of visual and parietal systems in primate spatial cognition (e.g., Rolls & Wirth, 2018; Martínez-Trujillo, 2025; Bottini et al., 2026), it remains unclear whether the human entorhinal cortex implements spatial maps in the same format. Third, in fMRI, the entorhinal cortex is particularly vulnerable to susceptibility-related signal loss and distortion because of nearby air– and bone–tissue interfaces, reducing sensitivity to representational effects relative to the hippocampus (Olman et al., 2009). Finally, entorhinal representations may not be map-like. For example, predictive-map accounts propose that entorhinal representations are graph-like, arising from the eigenstructure of experienced transitions, with different eigenmodes capturing graph structure at different scales (Stachenfeld et al., 2017). Such representations would not necessarily be captured by either Euclidean distance, which indexes map-like representations, or shortest-path distance, which mainly reflects low-order eigenmodes.

In conclusion, our findings reveal dissociable neural representations of cognitive maps and graphs in the human brain. The IPS may represent a cognitive map of spatial layout from an external survey perspective, whereas the hippocampus, together with scene-selective regions, may represent a cognitive graph structured by experiential transitions from an internal route perspective.

## Acknowledgments

This work was supported by a Marie Skłodowska-Curie Actions Postdoctoral Fellowship funded by the European Union (grant agreement number: 101064591; Y.X.), the Max Planck Society (C.F.D.), and the Max Planck School of Cognition (M.A.B.H.). We are grateful to Kerstin Schumer for her invaluable administrative and coordination support for the project. We appreciate Elisabeth Murzik’s generous support in organizing and facilitating data collection. We especially thank Paul Brockmeyer, Florian Muell, and Frieda Gabke for their dedicated assistance with data collection. We greatly benefited from the expertise and support of the MRI staff at the Max Planck Institute for Human Cognitive and Brain Sciences with MRI sequence setup, scheduling MRI sessions, and MRI data collection. Finally, we thank Simone Viganò and Andrej Bicanski for many helpful discussions.

## Author contributions

Y.X.: Conceptualization, Methodology, Formal analysis, Investigation, Data curation, Writing – original draft, Writing – review & editing, Visualization, Project administration, Funding acquisition.

M.A.B.H.: Conceptualization, Investigation, Writing – review & editing.

R.B.: Conceptualization, Writing – review & editing, Supervision.

C.F.D.: Conceptualization, Resources, Writing – review & editing, Supervision, Funding acquisition.

## Methods

### Participants

Sixty-one native German speakers with no history of neurological or psychiatric disorders participated in the experiment (age: M ± SD = 26.0 ± 4.6 years; gender: 32 female, 28 male, and 1 diverse; handedness: 60 right-handed and 1 left-handed). The fMRI data from one participant during the memory recall task were incomplete due to technical issues; analyses of this task were conducted using data from the remaining 60 participants.

The current study employed a whole-brain analysis to identify brain regions representing cognitive maps and graphs. Because this approach does not rely on pre-specified regional hypotheses, a hypothesis-driven power analysis based on predefined regions of interest was not feasible for determining sample size. Instead, sample size was informed by empirical evidence on the replicability of task-based fMRI findings, indicating that approximately 60 participants are required to achieve moderate stability of significant whole-brain clusters, as indexed by cluster overlap (Jaccard Index > 0.4; Turner et al., 2018).

The study was approved by the ethics committee at the University of Leipzig (reference number: 129/23-ek). All participants provided written informed consent and received compensation for their time.

### Stimuli

#### Virtual environment

We began by designing a Euclidean graph comprising eight locations that served as the blueprint for the virtual environment (Figure 1E). Figure S1A illustrates the corresponding map and graph information. By deliberately omitting direct connections between certain nearby locations, we orthogonalized Euclidean distance and shortest-path distance across the 28 possible location pairs (Figure 1F; r = −0.099). To provide an intuitive illustration of this orthogonalization, we performed clustering analysis on the Euclidean map and community detection on the graph structure. K-means clustering of the Euclidean coordinates identified two spatial clusters that can be separated by boundaries aligned with the lower-left–to–upper-right axis of the environment, with the optimal number of clusters determined using the silhouette method (Lloyd, 1982; Rousseeuw, 1987; Figure S1B). In contrast, modularity-based community detection on the adjacency matrix identified three modules that can be separated by boundaries aligned with the upper-left–to–lower-right axis of the environment (Newman, 2006; Figure S1C). These orthogonal partitions further illustrate the dissociation between map and graph information in the designed Euclidean graph.

We then used the Unity editor (version 2019.4.40f1) to implement the virtual environment, which consisted of a skybox and a set of highways. The skybox was uniformly shaded blue, containing no distal landmarks, clouds or sun sources, thereby minimizing external cues that could influence the formation of the intended cognitive maps or graphs. Highways were generated using the Path Creator toolbox (https://github.com/SebLague/Path-Creator), which creates paths using cubic Bézier curves. We defined the paths in the XZ plane (i.e., the ground plane), with anchor points placed at the Euclidean coordinates of the eight locations and connectivity determined by the adjacency matrix. The control mode of the Bézier curves was set to automatic, with the control-point spacing fixed at the minimum value, resulting in paths that closely approximate piecewise-linear segments with minimal curvature. Highways were constructed by extruding a road mesh (4 m wide and 0.01 m thick) along the defined paths. Invisible boundary walls were created using the same path-based extrusion procedure along both sides and at the termini of the highways; these invisible walls constrained participants to travel along the connected paths without obstructing the perception of spatial distances, thus preserving an open-maze environment.

#### Monuments

Each of the eight locations was marked by a unique sci-fi–style monument purchased from the Unity Asset Store (https://assetstore.unity.com/packages/3d/props/monuments-and-obelisks-110473). These eight monuments took the form of a variety of simple three-dimensional geometric shapes (Figure S2A). To standardize their physical scale, each monument was resized so that its longest dimension measured 4 m, and a 4 m × 4 m × 4 m box collider was assigned to each monument. All monuments were suspended above the ground, with their centres of mass positioned 3 m above the ground plane. The suspension configuration allowed the monuments to rotate continuously around the vertical axis at an angular speed of 120°/s, ensuring that their specific orientations did not influence the formation of the intended cognitive maps and graphs. To control for the effects of monument visual appearance on location representations, the eight monuments were assigned to eight locations in a Latin-square design across participants, achieving complete counterbalancing for most participants (56 out of 61).

Stimuli for the fMRI tasks were generated by capturing images of each monument from multiple viewing angles, with the camera positioned at the same height as the monument’s centre of mass and at a fixed distance of 5 m from it. For the standard condition, images were obtained from the eight learned monuments by rotating each monument in 40° increments over a full 360° rotation, yielding nine images per monument. For the oddball condition, images were obtained from an additional set of eight visually similar monuments drawn from the same Unity asset set but not previously learned by participants. Only a single view per monument was used in this condition (Figure S2B).

#### Categorical images

Images for the fMRI functional localizer to define the category-selective brain regions were obtained from a previous study (Josephs & Konkle, 2020) and downloaded from the associated online repository (https://osf.io/g9aj5/). From the folder “Stimuli_E1_StandardLocalizer,” we selected stimuli from three categories—scenes, faces, and objects—each comprising 40 images. Scene images consisted of indoor and outdoor views of navigable-scale environments. Face images were cropped from the chin to the top of the head and depicted individuals of varying ages and genders with diverse facial expressions. Object images depicted single objects on a white background. Object images were further scrambled by dividing them into 25 × 25 grids and randomly rearranging the grid elements, creating a fourth category of 40 images, referred to as scrambled images.

### Procedures

This study was conducted over two successive days. On the first day, participants learned the virtual environment outside the scanner through three learning stages, with behavioural tests administered after selected stages. On the second day, participants briefly revisited the virtual environment outside the scanner and then recalled information about each location in the cognitive map and graph during MRI data acquisition.

#### Virtual environment learning

The learning procedure was designed to allow participants to acquire both map and graph knowledge of the virtual environment. It consisted of four stages (Table S1). The first three stages were completed on the first day, and the fourth stage was conducted on the second day immediately before MRI scanning. The learning task was implemented and controlled using the OpenMaze toolbox (Alsbury-Nealy et al., 2022; https://openmaze.ca/). Across all four learning stages, participants used a computer keyboard to navigate the virtual environment as a 2-m-tall agent. The up and down arrow keys moved the agent forward and backward at a speed of 15 m/s, while the left and right arrow keys rotated the agent left and right at an angular speed of 60°/s (translation and rotation could not be performed simultaneously). Participants viewed the environment from a first-person, ground-level perspective. A third-person, survey perspective was not employed to preserve naturalistic navigation from a route perspective and avoid providing direct access to the overall spatial layout. To minimize directional carryover effects, participants’ initial facing direction was randomized on each trial, and a full 360° viewpoint rotation over 6 s was applied before navigation as a disorientation procedure.

The first stage was a familiarization stage. During this stage, all monuments were visible but inactive (see below). Participants were instructed to freely navigate the virtual environment for a fixed duration of 5 min in a single trial. The purpose of this stage was to provide participants with an overall impression of the environment. Navigation began from one of the midpoint locations along the highway segments (edges). There were eight edges, yielding eight possible starting positions, which were counterbalanced across participants in most cases (56 out of 61).

The second stage was the main learning stage. Participants started each trial from the midpoint location of one of the highway segments and were instructed to navigate to a target monument indicated at the top centre of the screen. The target monument and two additional monuments serving as competitors were invisible, with only the target monument being active: when the agent contacted the box collider surrounding the target monument, participants heard a coin sound, and the next trial began automatically. The remaining five monuments were visible but inactive, serving as visual cues for learning. With eight possible starting positions and eight possible goal locations, this stage comprised a total of 64 trials. To ensure balanced visual exposure, each monument was visible for an equal number of trials (40 trials per monument; 5 visible monuments × 64 trials ÷ 8 monuments). The trial order was randomized, and participants were allowed to take a break after 32 trials.

The third stage was a wrap-up stage. During this stage, all monuments were once again visible but inactive. Participants were instructed to freely navigate the virtual environment to obtain an overall view of the spatial layout of all monument locations and their connectivity. The settings for this stage were identical to those of the first learning stage, except that there was no time limit. Participants completed the stage when they felt they had acquired sufficient knowledge of the virtual environment.

The fourth stage was a warm-up stage conducted immediately before MRI scanning on the second day. This stage was identical to the third learning stage. Its purpose was to provide participants with a final opportunity to revisit the virtual environment, refresh what they had learned the previous day, and prepare for the tasks during scanning.

#### Behavioural testing

The behavioural tests were designed to monitor learning progression and ensure that participants acquired both map and graph knowledge of the virtual environment prior to MRI scanning. Assessments were conducted after Stages 1 and 3 (Figure 2B; Table S1). At both time points, participants completed a pairwise rating task evaluating Euclidean distance, shortest-path distance, and shape dissimilarity between monuments. Following Stage 3, participants additionally completed two tasks: a spatial arrangement task and a shortest-path selection task. All behavioural tests were implemented using PsychoPy (version 2022.2.4).

In the pairwise rating task, participants viewed images of two monuments presented side by side on the screen and used a horizontal slider to rate specific properties of the pair. In the Euclidean distance condition, participants rated the straight-line distance between the two monuments in the environment, regardless of whether a direct path connected them. In the shortest-path distance condition, participants rated the travel distance from the left monument to the right monument along the shortest possible route. In the shape dissimilarity condition, participants rated the dissimilarity between the monuments’ three-dimensional shapes. The right end of the slider corresponded to longer distances or greater dissimilarity. Each monument pair was presented twice per condition, with the monuments appearing in both possible left–right positions on the screen. There were 28 possible monument pairs, resulting in 56 trials per condition. Trial order was randomized within each condition. Slider responses were transformed to a continuous scale ranging from 0 to 7, and the two ratings obtained for each pair were averaged to produce the final rating.

In the spatial arrangement task, participants viewed images of eight monuments arranged around a circular arena on the screen and used the mouse to drag the icons into the arena so that their relative positions on the screen reflected the monuments’ locations in the environment.

In the shortest-path selection task, participants viewed images of two monuments presented side by side at the top of the screen and were instructed to imagine traveling from the left monument to the right monument along the shortest possible path. At the bottom of the screen, two additional monument images were presented as response options. Participants selected which monument they would pass through along the shortest path. Each monument pair at the top of the screen was presented twice, with the monuments appearing in both possible left–right positions. There were 20 possible monument pairs (i.e., the 28 total pairs excluding those that were directly connected), resulting in 40 trials. Trial order was randomized. On each trial, one monument located along the shortest path was randomly selected as the correct option, and another monument not on the shortest path was randomly selected as the distractor. The left–right positions of the correct option and distractor were also randomized.

#### MRI scanning

The MRI scanning session consisted of two runs of a functional localizer, two runs of a memory recall task, a T1-weighted structural scan, and six runs of a rating task, conducted sequentially. The rating task addressed a separate research question and is not analysed in the present study; it is the subject of a separate report. All fMRI stimuli were presented on a grey background.

The functional localizer was designed to identify category-selective brain regions using a block design with four conditions: scenes, faces, objects, and scrambled images. Within each run, condition order was counterbalanced within participants using a Latin square design, with each condition presented four times, resulting in 16 blocks per run. Within each block, participants maintained fixation on a centrally presented purple dot throughout the block while viewing images from a single category. Twenty unique images were selected from a pool of 40 images per category and presented in random order within each block. Across the four blocks per condition within a run, each image was presented twice. To maintain attention, one image from the first half of the block and one from the second half were randomly selected and immediately repeated (one-back targets), resulting in 22 trials per block. Participants performed a one-back task, responding as quickly as possible by pressing a button whenever an image was identical to the one immediately preceding it. Each image was displayed for 500 ms followed by a 500 ms interstimulus interval (trial duration = 1 s), yielding a block duration of 22 s. Each run began and ended with a 5 s blank interval, and blocks were separated by 5 s blank intervals.

The memory recall task was designed to guide participants to recall the map and graph knowledge of eight locations, cued by images of the corresponding monuments. In each run, the eight learned monuments were presented nine times from different viewing angles, resulting in 72 standard trials. To ensure that participants remained attentive throughout the experiment, an oddball condition was included: eight visually similar but previously unseen monuments were each presented once, yielding eight oddball trials and a total of 80 trials per run. All trials were pooled and presented in randomized order. During standard trials, participants were instructed to recall each monument’s position within the environment—both its location and its connections to other monuments—and to press a button with their right index finger once this information came to mind. During oddball trials, they pressed a different button using their right middle finger. Each run began with a 5.5 s blank screen and ended with a 4.5 s blank screen. Each trial started with a 500 ms fixation cross, followed by a 500 ms image presentation, and concluded with a 4 s blank screen.

### Behavioural analysis

#### Pairwise rating task during pre-scanning learning

To assess acquisition of cognitive map and graph knowledge, we computed Spearman correlations between participants’ pairwise distance ratings and the corresponding ground-truth distances, separately for Euclidean distance and shortest-path distance. Following the Stage 3 assessment, participants repeated Stage 3 if either correlation was below 0.8 or if the correlation between their Euclidean-distance and shortest-path-distance ratings exceeded 0.25.

To construct the shape-dissimilarity matrix for the subsequent RSA, we first assessed the consistency of participants’ shape-dissimilarity ratings using leave-one-participant-out Pearson correlations between each participant’s ratings and the mean ratings of all remaining participants. The resulting correlation coefficients were Fisher *z*-transformed, and outliers were identified as values below the first quartile by more than 1.5 times the interquartile range. One participant was identified as an outlier at both Stages 1 and 3 and was excluded. The shape-dissimilarity matrix used in the subsequent RSA was then calculated by averaging the remaining participants’ shape-dissimilarity ratings at Stage 3.

#### Spatial arrangement task during pre-scanning learning

We used the *mdscale* function in MATLAB R2025b to perform nonmetric multidimensional scaling (Kruskal, 1964) on the participant-averaged pairwise Euclidean distance matrix. Because participants may have used different spatial scales when arranging the icons on the screen, Euclidean distances were z-scored across location pairs within each participant before averaging across participants.

#### Shortest-path selection task during pre-scanning learning

Accuracy was calculated as the proportion of correctly selected responses.

#### Memory recall task during scanning

Accuracy was quantified as oddball detection sensitivity using *d’*, calculated as the difference between the z-transformed hit rate and false-alarm rate (Stanislaw & Todorov, 1999). Here, hits were defined as correct responses to oddball trials, and false alarms were defined as oddball responses on standard trials. To avoid infinite values arising from extreme hit or false-alarm rates, we applied the loglinear correction by adding 0.5 to the number of hits and false alarms and adding 1 to the number of oddball and standard trials before computing the rates (Hautus, 1995).

Reaction times (RTs) of the correct trials were analysed using JASP 0.19.3 with a linear mixed-effects model in which trial type (standard vs. oddball trials) was included as a fixed effect and by-participant random intercepts and slopes for trial type were included as random effects, specified as *RT* ∼ Trial Type + (1 + Trial Type ∣ Participant).

### MRI acquisition

MRI data were acquired using a MAGNETOM Skyra Fit 3T MR scanner (Siemens) with a 32-channel head-neck coil at the Max Planck Institute for Human Cognitive and Brain Sciences. Functional images were acquired using a simultaneous multislice gradient-echo echo-planar imaging (EPI) sequence, with anterior-to-posterior phase encoding and the scanning plane parallel to the long axis of the hippocampus: repetition time (TR) = 1000 ms, echo time (TE) = 22 ms, flip angle (FA) = 62°, field of view (FOV) = 204 mm × 204 mm, matrix size = 82 × 82, 65 axial slices, slice thickness = 2.5 mm, gap = 0.25 mm, voxel size = 2.49 × 2.49 × 2.75 mm, and multiband factor = 5.

Pairs of spin-echo EPI images with reversed phase-encoding directions were acquired between the two functional runs of each task to correct for geometric distortions in those runs. Each pair included one image acquired with anterior-to-posterior phase encoding and one image acquired with posterior-to-anterior phase encoding; the scanning plane was parallel to the long axis of the hippocampus: TR = 8000 ms, TE = 50 ms, FA = 90°, FOV = 204 mm × 204 mm, matrix size = 82 × 82, 65 axial slices, slice thickness = 2.5 mm, gap = 0.25 mm, and voxel size = 2.49 × 2.49 × 2.75 mm.

Three-dimensional T1-weighted images were acquired using the magnetization-prepared rapid gradient-echo sequence in the sagittal plane: TR = 2300 ms, TE = 5.28 ms, inversion time = 900 ms, FA = 9°, FOV = 240 mm × 256 mm, matrix size = 240 × 256, 176 continuous sagittal slices, slice thickness = 1 mm, and voxel size = 1 × 1 × 1 mm.

### MRI preprocessing

We performed MRI preprocessing using fMRIPrep 22.0.1 (Esteban et al., 2019), which is based on Nipype 1.8.4 (Gorgolewski et al., 2011). Detailed preprocessing steps are described in the Supplemental Methods using boilerplate text generated directly by fMRIPrep, which provides a clear and standardized description to improve reproducibility. For subsequent analyses, we used the functional images in MNI152NLin6Asym space generated using independent component analysis-based automatic removal of motion artifacts (ICA-AROMA; Pruim et al., 2015).

### Independent Resting-State fMRI Dataset

For the functional connectivity analysis, we used an independent, openly available resting-state fMRI dataset acquired at our institution: the Leipzig Study for Mind–Body–Emotion Interactions (LEMON) dataset. The data were obtained from the public repository (https://ftp.gwdg.de/pub/misc/MPI-Leipzig_Mind-Brain-Body-LEMON/MRI_MPILMBB_LEMON/). Detailed information on participant recruitment, MRI acquisition, and preprocessing is provided in the corresponding data descriptor (Babayan et al., 2019); a brief summary is presented below.

The dataset includes both younger and older adults. For the present analysis, we included only the younger group (N = 154; mean age = 25.1 ± 3.1 years; age range = 20–35 years; 45 females). Two participants were excluded: one because of a corrupted imaging file and one because resting-state fMRI data were unavailable. The final sample therefore comprised 152 participants.

The resting-state fMRI sequence had a repetition time of 1,400 ms and comprised 657 volumes. We used the preprocessed resting-state fMRI data provided with the dataset. Preprocessing included: (i) removal of the first five volumes; (ii) motion correction; (iii) distortion correction; (iv) rigid-body coregistration; (v) denoising; (vi) temporal band-pass filtering between 0.01 and 0.10 Hz; and (vii) spatial normalization to 2-mm MNI152 standard space.

### fMRI analysis

#### First-level fMRI analysis

The first-level fMRI analysis was performed using Statistical Parametric Mapping 12 (SPM 12). Individual-level general linear models (GLMs) were constructed separately for the univariate contrasts in the functional localizer, the univariate contrasts in the memory recall task, and the representational similarity analyses in the memory recall task. In all three GLMs, nuisance regressors extracted from fMRIPrep included mean time courses from conservative white-matter and cerebrospinal-fluid masks, their squared terms, their temporal derivatives, their squared temporal derivatives, and the Friston 24-parameter head-motion model (Friston et al., 1996). Run-specific constant regressors were also included in each GLM. A high-pass filter with a cutoff of 128 s was used to remove low-frequency noise and slow drifts.

The GLM for the univariate contrast analysis in the functional localizer included four conditions: scenes, faces, objects, and scrambled objects. For each condition, the onset and duration of the corresponding blocks were used to construct boxcar regressors, which were then convolved with the canonical hemodynamic response function (HRF). Scene-selective regions were identified by contrasting scenes with the average of the face and object conditions.

The GLMs for the memory recall task used a common event-modelling approach. For each trial, the onset was defined as the stimulus onset, and the duration was defined as the participant’s RT. These trial-wise boxcar regressors were convolved with the canonical HRF, using a variable-epoch approach to model trial-by-trial RT variability (Grinband et al., 2008). To further control for domain-general RT-related effects across conditions, all trials were pooled and modelled with an additional stick-function regressor at trial onset, parametrically modulated by mean-centred RT. This RT-modulated regressor was convolved with the canonical HRF and included in the GLM as a nuisance regressor.

For the univariate contrast analysis, the GLM included two conditions: standard trials and oddball trials. Contrasts were computed between standard and oddball trials, and the resulting contrast beta images were entered into the second-level analyses. For the representational similarity analysis (RSA), the GLM included nine conditions: the eight locations cued by the corresponding monuments and the oddball trials. Each location was contrasted against the implicit baseline, and the resulting T images, rather than beta images, were used in the subsequent RSA (Misaki et al., 2010).

#### Resting-state functional connectivity analysis

To characterize the functional networks associated with the anterior and posterior hippocampus, we performed seed-based resting-state functional connectivity (RSFC) analyses using the independent resting-state fMRI dataset described above. The seed regions were defined based on the univariate analysis of the main experiment. The anterior hippocampal seed comprised voxels showing greater activation in the standard than in the oddball condition, whereas the posterior hippocampal seed comprised voxels showing greater activation in the oddball than in the standard condition. Both seeds were defined using a cluster-forming threshold of p < 0.001 and a whole-brain cluster-level FWE– corrected threshold of p < 0.05. For each participant, the resting-state time series of all voxels within each seed were extracted and averaged to obtain a representative seed time series. The seed time series was correlated with the time series of every voxel in the brain using Pearson correlation. The resulting correlation coefficients were then Fisher z-transformed. This procedure yielded two whole-brain RSFC maps for each participant: one for the anterior hippocampal seed and one for the posterior hippocampal seed.

To directly compare the connectivity profiles of the two hippocampal regions, the posterior hippocampal RSFC map was subtracted from the anterior hippocampal RSFC map for each participant. The resulting individual contrast maps were spatially smoothed with an isotropic Gaussian kernel with a full-width at half-maximum of 8 mm before second-level analysis.

#### Searchlight-based representational similarity analysis

RSA was conducted across the eight locations (Kriegeskorte et al., 2008). We calculated the Euclidean distance between voxel-wise activity patterns for each pair of locations, yielding an 8 × 8 neural representational dissimilarity matrix (RDM). We then correlated the neural RDM with each model RDM—Euclidean distance, shortest-path distance, and shape dissimilarity—across the 28 unique location pairs using Pearson correlation. Because monument-to-location assignments varied across participants according to the Latin-square design, we reordered the shape-dissimilarity RDM separately for each participant to match that participant’s monument-to-location assignment before correlating it with the neural RDM. The resulting correlation coefficients were Fisher *z*-transformed.

Searchlight-based RSA was performed across the whole brain (Kriegeskorte et al., 2006). The searchlight was moved across all voxels within a whole-brain grey-matter mask, defined using the Harvard–Oxford cortical and subcortical structural atlases with a 25% probability threshold and including the bilateral cerebral cortex, caudate, putamen, pallidum, accumbens, hippocampus, and amygdala. Because there is no established optimal searchlight radius, we examined radii of 6, 8, and 10 mm. These analyses yielded very similar spatial patterns, with effects becoming stronger at larger radii. We therefore present the results obtained with the 10-mm radius. For each searchlight, the Fisher z-transformed Pearson correlation between the neural RDM and each model RDM was assigned to the centre voxel of the searchlight. The resulting searchlight maps were spatially smoothed with an isotropic Gaussian kernel with a full-width at half-maximum of 8 mm before second-level analysis.

To zoom in on the hippocampal–entorhinal system, we performed an anatomically restricted searchlight RSA encompassing the hippocampal formation and entorhinal cortex. We defined the search volume using bilateral probabilistic maps from the 207-region Jülich Brain Atlas v3.1 (Amunts et al., 2020, 2023), including hippocampal CA1, CA2, CA3, dentate gyrus (DG), hippocampal–amygdaloid transition area (HATA), subiculum (Subc), hippocampal transition area (Trs), and entorhinal cortex (Area EC). These regions are based on cytoarchitectonic probabilistic mappings of the human hippocampal region and entorhinal cortex (Amunts et al., 2005). The probabilistic maps were transformed from MNI152NLin2009cAsym to MNI152NLin6Asym space using Advanced Normalization Tools (ANTs) with linear interpolation and then combined into a single probability map by taking the voxel-wise maximum across all regions. The resulting map was thresholded at 25% probability to generate a binary hippocampal–entorhinal mask. Searchlight RSA was then repeated strictly within this mask using a 10-mm radius. Both searchlight centres and the voxels contributing to each neural RDM were constrained to lie within the anatomical mask. No additional spatial smoothing was applied to the resulting searchlight RSA maps before second-level analysis, thereby limiting further spatial blurring across adjacent hippocampal–entorhinal subregions and anatomical boundaries.

#### Second-level fMRI analysis

The second-level fMRI analyses were performed using the Permutation Analysis of Linear Models (PALM) toolbox. For the whole-brain analyses, statistical inference and multiple-comparison correction were restricted to a whole-brain grey-matter mask defined from the Harvard–Oxford cortical and subcortical structural atlases using a 25% probability threshold. The mask encompassed the bilateral cerebral cortex, caudate, putamen, pallidum, nucleus accumbens, hippocampus, and amygdala. For the anatomically restricted searchlight analysis of the hippocampal–entorhinal system, statistical inference and multiple-comparison correction were restricted to a predefined hippocampal–entorhinal mask constructed from cytoarchitectonic probabilistic maps from the Jülich Brain Atlas using a 25% probability threshold, as described above. The mask encompassed the hippocampal formation and entorhinal cortex. Null distributions were generated using 5,000 sign flips (Winkler et al., 2014). To obtain more precise estimates of small *p*-values, a generalized Pareto distribution was fitted to the tail of the permutation distribution for *p*-values below 0.10 (Knijnenburg et al., 2009; Winkler et al., 2016).

For the univariate contrasts from the functional localizer and memory-recall task, as well as the searchlight-based representational similarity analyses in the memory-recall task, family-wise error was controlled using cluster-level correction with a voxel-wise cluster-forming threshold of p < 0.001 and a cluster-level FWE-corrected threshold of p < 0.05 (Woo et al., 2014). For the RSFC analysis, this cluster-level procedure yielded spatially extensive clusters that were insufficiently anatomically specific. We therefore applied a more stringent voxel-wise threshold, controlling whole-brain FWE at p < 0.05.

To quantify the spatial correspondence between shortest-path-distance effects and scene-selective activation, we calculated the percentage of voxels within each significant shortest-path-distance cluster that fell within the thresholded scene-selective activation map.

#### Network correspondence analysis

We used the Network Correspondence Toolbox (Kong et al., 2025) to characterize the spatial correspondence between task-activation and RSFC maps with canonical functional brain networks. Specifically, we examined the network correspondence of the contrasts between the standard and oddball conditions and between the anterior and posterior hippocampal RSFC maps. The input images were statistical maps thresholded according to the thresholds applied in the corresponding second-level analyses. For the task analyses, we entered thresholded z-statistic maps contrasting standard and oddball conditions, obtained using a voxel-wise cluster-forming threshold of p < 0.001 and a whole-brain cluster-level FWE-corrected threshold of p < 0.05. For the RSFC analyses, we entered thresholded z-statistic maps contrasting anterior and posterior hippocampal connectivity, obtained using whole-brain voxel-wise FWE correction at p < 0.05.

The toolbox binarized the thresholded z-statistic maps and quantified their spatial correspondence with each network in the Yeo 17-network atlas using the Dice coefficient (Yeo et al., 2011). Statistical significance was assessed using spin-test permutations. For this analysis, the volumetric MNI-space maps were first projected onto the fsaverage6 cortical surface. The spatial pattern of each map was then randomly rotated 1,000 times while preserving its spatial structure. For each rotation and atlas network, the Dice coefficient was recalculated to generate a null distribution. The permutation p-value was calculated as the proportion of rotations yielding a Dice coefficient greater than that observed for the original map.

#### ROI-based representational similarity analysis

We conducted ROI-based RSA using the same procedure described in the *searchlight-based representational similarity analysis* section, with neural RDMs constructed from voxel-wise activity patterns within each ROI. The definitions of the ROIs and analysis-specific details are described below.

To characterize the representational profiles of the regions identified in the whole-brain searchlight analysis, we performed post-hoc ROI-based RSA in the right intraparietal sulcus, right superior lateral occipital cortex, right posteromedial region, and left hippocampal/parahippocampal region. The ROIs comprised voxels within the corresponding significant searchlight clusters, identified using a voxel-wise cluster-forming threshold of *p* < 0.001 and a cluster-level FWE-corrected threshold of *p* < 0.05. Because the ROIs were defined from the searchlight analysis itself, we did not treat these analyses as independent tests of the searchlight effects. Rather, they served as follow-up analyses of the searchlight results (Etzel et al., 2013), assessing whether the voxels within the identified clusters showed the corresponding representational effects and characterizing their specificity. Specifically, we tested whether the right intraparietal sulcus showed evidence for shortest-path distance and whether the three regions identified by the shortest-path searchlight showed evidence for Euclidean distance. Participant-level Fisher *z*-transformed correlation coefficients were tested against zero using one-sample *t*-tests. To further assess whether the relative representation of Euclidean and shortest-path distance differed across regions, we conducted a 4 × 2 repeated-measures ANOVA with Region and Representation as within-participant factors. To visualize the representational relationships underlying these effects, we additionally plotted neural distance against the corresponding model distance for each of the 28 location pairs. Neural distances were *z*-standardized across the 28 pairs within each participant and then averaged across participants. Mean neural distance was plotted against Euclidean distance for the right intraparietal sulcus and against shortest-path distance for the right superior lateral occipital cortex, right posteromedial region, and left hippocampal/parahippocampal region. Linear regression lines with 95% confidence intervals were included for descriptive visualization only.

As a post-hoc robustness analysis, we further examined whether the representational effects within the searchlight-defined ROIs remained after accounting for shared variance among the model RDMs. Pearson partial correlations were computed separately for Euclidean distance, shortest-path distance, and shape dissimilarity, with each model tested while controlling for the other two model RDMs. For each participant, the resulting partial correlation coefficients were Fisher *z*-transformed and then tested against zero across participants using one-sample *t*-tests. *P*-values were Holm-adjusted across the three models within each ROI.

We additionally examined representations within independently constrained, functionally localized scene-selective regions, including the parahippocampal place area (PPA), retrosplenial cortex (RSC), and occipital place area (OPA), separately in the left and right hemispheres. Scene-selective activation was defined using the functional-localizer contrast of scenes > faces/objects, with a voxel-wise cluster-forming threshold of *p* < 0.001 and a cluster-level FWE-corrected threshold of *p* < 0.05. Because the resulting scene-selective activation was spatially connected across these regions, we used independently derived scene-selective parcels to constrain the functional localizer and delineate the conventional PPA, RSC, and OPA (Julian et al., 2012; https://web.mit.edu/bcs/nklab/GSS.shtml). The original parcels, provided in SPM5 MNI template space, were nonlinearly transformed to MNI152NLin6Asym space using ANTs, with nearest-neighbour interpolation applied to preserve the discrete parcel labels. The transverse occipital sulcus (TOS) parcel was used as the parcel corresponding to the OPA. Each transformed parcel was then intersected with the corresponding thresholded scene-selective activation to define the final ROI. Within each ROI and hemisphere, neural RDMs were correlated with the Euclidean-distance, shortest-path-distance, and shape-dissimilarity model RDMs using Pearson correlation, and the resulting coefficients were Fisher *z*-transformed. Participant-level Fisher *z*-transformed coefficients were tested against zero using one-sample *t*-tests. To control for multiple comparisons, *p*-values were Holm-adjusted across the six tests comprising the three model RDMs × two hemispheres within each ROI.

Finally, to specifically examine representations in the entorhinal cortex within the hippocampal– entorhinal system, we performed ROI-based RSA separately in the left and right entorhinal cortex. The ROIs were defined from the cytoarchitectonic Area EC probabilistic maps of the Jülich Brain Atlas described above; following transformation to MNI152NLin6Asym space, the left and right maps were thresholded separately at 25% probability to generate binary entorhinal cortex masks. Within each hemisphere, neural RDMs were correlated with the Euclidean-distance, shortest-path-distance, and shape-dissimilarity model RDMs using Pearson correlation, and the resulting coefficients were Fisher *z*-transformed. Participant-level Fisher *z*-transformed coefficients were tested against zero using one-sample *t*-tests.

#### Brain map visualization

To provide a global overview of the whole-brain results, we visualized thresholded z-statistic maps of significant effects and scene-selective regions on the cortical surface using Connectome Workbench version 2.2.1 (https://www.humanconnectome.org/software/connectome-workbench). The volumetric z-statistic maps were projected onto the fsLR surface using the neuromaps toolbox (Markello et al., 2022). The MNI152NLin6Asym-transformed scene-selective volumetric parcel masks were converted to label volumes and mapped onto the corresponding 32k fsLR midthickness surface using ribbon-constrained volume-to-surface mapping between the white and pial surfaces in Connectome Workbench. The resulting surface labels were converted to border files using *wb_command-label-to-border* and displayed on the cortical surface as needed. The surface maps were displayed on an inflated cortical surface overlaid with the group-average sulcal-depth map derived from 1,096 young adults in the Human Connectome Project dataset (https://balsa.wustl.edu/reference/pkXDZ).

To visualize subcortical structures, particularly the hippocampus, the same thresholded z-statistic maps were additionally displayed in volumetric space using MRIcroGL. The maps were overlaid on the 0.5-mm isotropic tpl-MNI152NLin6Asym_res-03_T1w.nii.gz anatomical template obtained from TemplateFlow (https://www.templateflow.org/).

## References

Alexander, A. S., & Nitz, D. A. (2015). Retrosplenial cortex maps the conjunction of internal and external spaces. Nature Neuroscience, 18(8), 1143–1151.

Alexander, A. S., & Nitz, D. A. (2017). Spatially periodic activation patterns of retrosplenial cortex encode route sub-spaces and distance traveled. Current Biology, 27(11), 1551–1560.

Alexander, A. S., Place, R., Starrett, M. J., Chrastil, E. R., & Nitz, D. A. (2023). Rethinking retrosplenial cortex: Perspectives and predictions. Neuron, 111(2), 150–175.

Alsbury-Nealy, K., Wang, H., Howarth, C., Gordienko, A., Schlichting, M. L., & Duncan, K. D. (2022). OpenMaze: An open-source toolbox for creating virtual navigation experiments. Behavior Research Methods, 54(3), 1374–1387.

Amunts, K., Kedo, O., Kindler, M., Pieperhoff, P., Mohlberg, H., Shah, N. J., … & Zilles, K. (2005). Cytoarchitectonic mapping of the human amygdala, hippocampal region and entorhinal cortex: intersubject variability and probability maps. Anatomy and Embryology, 210(5), 343–352.

Amunts, K., Mohlberg, H., Bludau, S., & Zilles, K. (2020). Julich-Brain: A 3D probabilistic atlas of the human brain’s cytoarchitecture. Science, 369(6506), 988–992.

Amunts, K., Mohlberg, H., Bludau, S., Caspers, S., Lewis, L. B., Eickhoff, S. B., & Pieperhoff, P. (2023). *Julich-Brain Atlas, cytoarchitectonic maps* (Version 3.1) [Data set]. EBRAINS. 10.25493/KNSN-XB4

Angeli, P. A., DiNicola, L. M., Saadon-Grosman, N., Eldaief, M. C., & Buckner, R. L. (2025). Specialization of the human hippocampal long axis revisited. Proceedings of the National Academy of Sciences, 122(3), e2422083122.

Babayan, A., Erbey, M., Kumral, D., Reinelt, J. D., Reiter, A. M., Röbbig, J., … & Villringer, A. (2019). A mind-brain-body dataset of MRI, EEG, cognition, emotion, and peripheral physiology in young and old adults. Scientific Data, 6(1), 180308.

Baldassano, C., Esteva, A., Fei-Fei, L., & Beck, D. M. (2016). Two distinct scene-processing networks connecting vision and memory. eNeuro, 3(5), ENEURO.0178-16.2016.

Bálint, R. (1909). Seelenlähmung des “schauens”, optische ataxie, räumliche störung der aufmerksamkeit. Monatsschr Psychiatr Neurol, 25(1), 51–81.

Behrens, T. E., Muller, T. H., Whittington, J. C., Mark, S., Baram, A. B., Stachenfeld, K. L., & Kurth-Nelson, Z. (2018). What is a cognitive map? Organizing knowledge for flexible behavior. Neuron, 100(2), 490–509.

Bellmund, J. L., Gärdenfors, P., Moser, E. I., & Doeller, C. F. (2018). Navigating cognition: Spatial codes for human thinking. Science, 362(6415), eaat6766.

Bisley, J. W., & Goldberg, M. E. (2010). Attention, intention, and priority in the parietal lobe. Annual review of neuroscience, 33(1), 1–21.

Bittner, K. C., Milstein, A. D., Grienberger, C., Romani, S., & Magee, J. C. (2017). Behavioral time scale synaptic plasticity underlies CA1 place fields. Science, 357(6355), 1033–1036.

Bonner, M. F., & Epstein, R. A. (2017). Coding of navigational affordances in the human visual system. Proceedings of the National Academy of Sciences, 114(18), 4793–4798.

Boroditsky, L. (2000). Metaphoric structuring: Understanding time through spatial metaphors. Cognition, 75(1), 1–28.

Bottini, R., Viganò, S., Xu, Y., van Dijck, J.-P., Abrahamse, E., & Pezzulo, G. (2026). Actionable cognitive maps for spatial and conceptual thinking. PsyArXiv.

Burak, Y., & Fiete, I. R. (2009). Accurate path integration in continuous attractor network models of grid cells. PLoS Computational Biology, 5(2), e1000291.

Buzsáki, G., & Tingley, D. (2018). Space and time: the hippocampus as a sequence generator. Trends in Cognitive Sciences, 22(10), 853–869.

Chechlacz, M., & Humphreys, G. W. (2014). The enigma of Bálint’s syndrome: neural substrates and cognitive deficits. Frontiers in Human Neuroscience, 8, 123.

Corbetta, M., & Shulman, G. L. (2002). Control of goal-directed and stimulus-driven attention in the brain. Nature Reviews Neuroscience, 3(3), 201–215

Corbetta, M., Kincade, J. M., & Shulman, G. L. (2002). Neural systems for visual orienting and their relationships to spatial working memory. Journal of Cognitive Neuroscience, 14(3), 508–523.

Dede, A. J., Frascino, J. C., Wixted, J. T., & Squire, L. R. (2016). Learning and remembering real-world events after medial temporal lobe damage. Proceedings of the National Academy of Sciences, 113(47), 13480–13485.

Dilks, D. D., Julian, J. B., Paunov, A. M., & Kanwisher, N. (2013). The occipital place area is causally and selectively involved in scene perception. The Journal of Neuroscience, 33(4), 1331–1336.

Doeller, C. F., Barry, C., & Burgess, N. (2010). Evidence for grid cells in a human memory network. Nature, 463(7281), 657–661.

Eichenbaum, H. (2014). Time cells in the hippocampus: a new dimension for mapping memories. Nature Reviews Neuroscience, 15(11), 732–744.

Eichenbaum, H., & Cohen, N. J. (2001). From conditioning to conscious recollection: Memory systems of the brain. Oxford University Press.

Eichenbaum, H., Dudchenko, P., Wood, E., Shapiro, M., & Tanila, H. (1999). The hippocampus, memory, and place cells: is it spatial memory or a memory space? Neuron, 23(2), 209–226.

Epstein, R. A., & Baker, C. I. (2019). Scene perception in the human brain. Annual Review of Vision Science, 5(1), 373–397.

Epstein, R. A., Parker, W. E., & Feiler, A. M. (2007). Where am I now? Distinct roles for parahippocampal and retrosplenial cortices in place recognition. Journal of Neuroscience, 27(23), 6141–6149.

Epstein, R., & Kanwisher, N. (1998). A cortical representation of the local visual environment. Nature, 392(6676), 598–601.

Esteban, O., Markiewicz, C. J., Blair, R. W., Moodie, C. A., Isik, A. I., Erramuzpe, A., … & Gorgolewski, K. J. (2019). fMRIPrep: a robust preprocessing pipeline for functional MRI. Nature Methods, 16(1), 111–116.

Etzel, J. A., Zacks, J. M., & Braver, T. S. (2013). Searchlight analysis: promise, pitfalls, and potential. NeuroImage, 78, 261–269.

Friston, K. J., Williams, S., Howard, R., Frackowiak, R. S., & Turner, R. (1996). Movement-related effects in fMRI time-series. Magnetic Resonance in Medicine, 35(3), 346–355.

Fyhn, M., Hafting, T., Treves, A., Moser, M. B., & Moser, E. I. (2007). Hippocampal remapping and grid realignment in entorhinal cortex. Nature, 446(7132), 190–194.

Gardner, R. J., Hermansen, E., Pachitariu, M., Burak, Y., Baas, N. A., Dunn, B. A., … & Moser, E. I. (2022). Toroidal topology of population activity in grid cells. Nature, 602(7895), 123–128.

Garvert, M. M., Dolan, R. J., & Behrens, T. E. (2017). A map of abstract relational knowledge in the human hippocampal–entorhinal cortex. eLife, 6, e17086.

George, D., Rikhye, R. V., Gothoskar, N., Guntupalli, J. S., Dedieu, A., & Lázaro-Gredilla, M. (2021). Clone-structured graph representations enable flexible learning and vicarious evaluation of cognitive maps. Nature Communications, 12(1), 2392.

Goodale, M. A., & Milner, A. D. (1992). Separate visual pathways for perception and action. Trends in Neurosciences, 15(1), 20–25.

Gorgolewski, K., Burns, C. D., Madison, C., Clark, D., Halchenko, Y. O., Waskom, M. L., & Ghosh, S. S. (2011). Nipype: a flexible, lightweight and extensible neuroimaging data processing framework in python. Frontiers in Neuroinformatics, 5, 13.

Gottlieb, J. P., Kusunoki, M., & Goldberg, M. E. (1998). The representation of visual salience in monkey parietal cortex. Nature, 391(6666), 481–484.

Grinband, J., Wager, T. D., Lindquist, M., Ferrera, V. P., & Hirsch, J. (2008). Detection of time-varying signals in event-related fMRI designs. NeuroImage, 43(3), 509–520.

Hafting, T., Fyhn, M., Molden, S., Moser, M. B., & Moser, E. I. (2005). Microstructure of a spatial map in the entorhinal cortex. Nature, 436(7052), 801–806.

Hautus, M. J. (1995). Corrections for extreme proportions and their biasing effects on estimated values of d′. Behavior Research Methods, Instruments, & Computers, 27(1), 46–51.

Howard, L. R., Javadi, A. H., Yu, Y., Mill, R. D., Morrison, L. C., Knight, R., … & Spiers, H. J. (2014). The hippocampus and entorhinal cortex encode the path and Euclidean distances to goals during navigation. Current Biology, 24(12), 1331–1340.

Jacobs, J., Weidemann, C. T., Miller, J. F., Solway, A., Burke, J. F., Wei, X. X., … & Kahana, M. J. (2013). Direct recordings of grid-like neuronal activity in human spatial navigation. Nature Neuroscience, 16(9), 1188–1190.

Josephs, E. L., & Konkle, T. (2020). Large-scale dissociations between views of objects, scenes, and reachable-scale environments in visual cortex. Proceedings of the National Academy of Sciences, 117(47), 29354–29362.

Julian, J. B., Fedorenko, E., Webster, J., & Kanwisher, N. (2012). An algorithmic method for functionally defining regions of interest in the ventral visual pathway. NeuroImage, 60(4), 2357–2364.

Kim, H. (2021). Imaging recollection, familiarity, and novelty in the frontoparietal control and default mode networks and the anterior-posterior medial temporal lobe: an integrated view and meta-analysis. Neuroscience & Biobehavioral Reviews, 126, 491–508.

Knijnenburg, T. A., Wessels, L. F., Reinders, M. J., & Shmulevich, I. (2009). Fewer permutations, more accurate P-values. Bioinformatics, 25(12), i161–i168.

Kong, R., Spreng, R. N., Xue, A., Betzel, R. F., Cohen, J. R., Damoiseaux, J. S., … & Uddin, L. Q. (2025). A network correspondence toolbox for quantitative evaluation of novel neuroimaging results. Nature Communications, 16(1), 2930.

Kriegeskorte, N., Goebel, R., & Bandettini, P. (2006). Information-based functional brain mapping. Proceedings of the National Academy of Sciences, 103(10), 3863–3868.

Kriegeskorte, N., Mur, M., & Bandettini, P. A. (2008). Representational similarity analysis-connecting the branches of systems neuroscience. Frontiers in Systems Neuroscience, 2, 4.

Kruskal, J. B. (1964). Nonmetric multidimensional scaling: a numerical method. Psychometrika, 29(2), 115–129.

Lakoff, G., & Johnson, M. (1980). Metaphors we live by. University of Chicago Press.

Li, Y., Briguglio, J. J., Romani, S., & Magee, J. C. (2024). Mechanisms of memory-supporting neuronal dynamics in hippocampal area CA3. Cell, 187(24), 6804–6819.

Lisman, J., Buzsáki, G., Eichenbaum, H., Nadel, L., Ranganath, C., & Redish, A. D. (2017). Viewpoints: how the hippocampus contributes to memory, navigation and cognition. Nature Neuroscience, 20(11), 1434–1447.

Lloyd, S. (1982). Least squares quantization in PCM. IEEE Transactions on Information Theory, 28(2), 129–137.

Markello, R. D., Hansen, J. Y., Liu, Z. Q., Bazinet, V., Shafiei, G., Suárez, L. E., … & Misic, B. (2022). Neuromaps: structural and functional interpretation of brain maps. Nature Methods, 19(11), 1472–1479.

Martínez-Trujillo, J. (2025). Why do primates have view cells instead of place cells? Trends in Cognitive Sciences, 29(3), 226–229.

McNaughton, B. L., Battaglia, F. P., Jensen, O., Moser, E. I., & Moser, M. B. (2006). Path integration and the neural basis of the “cognitive map”. Nature Reviews Neuroscience, 7(8), 663–678.

Misaki, M., Kim, Y., Bandettini, P. A., & Kriegeskorte, N. (2010). Comparison of multivariate classifiers and response normalizations for pattern-information fMRI. NeuroImage, 53(1), 103–118.

Mishkin, M., Ungerleider, L. G., & Macko, K. A. (1983). Object vision and spatial vision: two cortical pathways. Trends in Neurosciences, 6, 414–417.

Morgan, L. K., MacEvoy, S. P., Aguirre, G. K., & Epstein, R. A. (2011). Distances between real-world locations are represented in the human hippocampus. The Journal of Neuroscience, 31(4), 1238–1245.

Muller, R. U., Stead, M., & Pach, J. (1996). The hippocampus as a cognitive graph. The Journal of General Physiology, 107(6), 663–694.

Newman, M. E. (2006). Modularity and community structure in networks. Proceedings of the National Academy of Sciences, 103(23), 8577–8582.

O’Keefe, J., & Dostrovsky, J. (1971). The hippocampus as a spatial map: preliminary evidence from unit activity in the freely-moving rat. Brain Research, 34(1), 171–175.

O’Keefe, J., & Nadel, L. (1978). The hippocampus as a cognitive map. Oxford University Press.

Olman, C. A., Davachi, L., & Inati, S. (2009). Distortion and signal loss in medial temporal lobe. PLOS One, 4(12), e8160.

Ottink, L., de Haas, N., & Doeller, C. F. (2025). Integration of Euclidean and path distances in hippocampal maps. Scientific Reports, 15(1), 7104.

Peer, M., Brunec, I. K., Newcombe, N. S., & Epstein, R. A. (2021). Structuring knowledge with cognitive maps and cognitive graphs. Trends in Cognitive Sciences, 25(1), 37–54.

Peer, M., Nadar, C., & Epstein, R. A. (2024). The format of the cognitive map depends on the structure of the environment. Journal of Experimental Psychology: General, 153(1), 224–240.

Pruim, R. H., Mennes, M., van Rooij, D., Llera, A., Buitelaar, J. K., & Beckmann, C. F. (2015). ICA-AROMA: A robust ICA-based strategy for removing motion artifacts from fMRI data. NeuroImage, 112, 267–277.

Quent, J. A., Song, L., Liang, X., Su, Y., Yu, W., Wang, H., & Vatansever, D. (2026). Graded encoding of spatial novelty scales in the human brain. Nature Communications, 17(1), 303.

Rizzo, M., & Vecera, S. P. (2002). Psychoanatomical substrates of Balint’s syndrome. *Journal of Neurology*, Neurosurgery & Psychiatry, 72(2), 162–178.

Rolls, E. T., & Wirth, S. (2018). Spatial representations in the primate hippocampus, and their functions in memory and navigation. Progress in Neurobiology, 171, 90–113.

Rousseeuw, P. J. (1987). Silhouettes: a graphical aid to the interpretation and validation of cluster analysis. Journal of Computational and Applied Mathematics, 20, 53–65.

Schapiro, A. C., Turk-Browne, N. B., Norman, K. A., & Botvinick, M. M. (2016). Statistical learning of temporal community structure in the hippocampus. Hippocampus, 26(1), 3–8.

Scoville, W. B., & Milner, B. (1957). Loss of recent memory after bilateral hippocampal lesions. *Journal of Neurology*, Neurosurgery, and Psychiatry, 20(1), 11–21.

Sereno, M. I., Pitzalis, S., & Martinez, A. (2001). Mapping of contralateral space in retinotopic coordinates by a parietal cortical area in humans. Science, 294(5545), 1350–1354.

Shrager, Y., Kirwan, C. B., & Squire, L. R. (2008). Neural basis of the cognitive map: Path integration does not require hippocampus or entorhinal cortex. Proceedings of the National Academy of Sciences, 105(33), 12034–12038.

Siegel, A. W., & White, S. H. (1975). The development of spatial representations of large-scale environments. In Advances in Child Development and Behavior (Vol. 10, pp. 9–55). JAI.

Silver, M. A., & Kastner, S. (2009). Topographic maps in human frontal and parietal cortex. Trends in Cognitive Sciences, 13(11), 488–495

Silver, M. A., Ress, D., & Heeger, D. J. (2005). Topographic maps of visual spatial attention in human parietal cortex. Journal of Neurophysiology, 94(2), 1358–1371.

Stachenfeld, K. L., Botvinick, M. M., & Gershman, S. J. (2017). The hippocampus as a predictive map. Nature Neuroscience, 20(11), 1643–1653.

Stanislaw, H., & Todorov, N. (1999). Calculation of signal detection theory measures. Behavior Research Methods, Instruments, & Computers, 31(1), 137–149.

Swisher, J. D., Halko, M. A., Merabet, L. B., McMains, S. A., & Somers, D. C. (2007). Visual topography of human intraparietal sulcus. The Journal of Neuroscience, 27(20), 5326–5337.

Tullo, M. G., Almgren, H., Van de Steen, F., Boccia, M., Bencivenga, F., & Galati, G. (2023). Preferential signal pathways during the perception and imagery of familiar scenes: An effective connectivity study. Human Brain Mapping, 44(10), 3954–3971.

Turner, B. O., Paul, E. J., Miller, M. B., & Barbey, A. K. (2018). Small sample sizes reduce the replicability of task-based fMRI studies. Communications Biology, 1(1), 62.

Warren, W. H. (2019). Non-euclidean navigation. Journal of Experimental Biology, 222(Suppl_1), jeb187971.

Winkler, A. M., Ridgway, G. R., Douaud, G., Nichols, T. E., & Smith, S. M. (2016). Faster permutation inference in brain imaging. NeuroImage, 141, 502–516.

Winkler, A. M., Ridgway, G. R., Webster, M. A., Smith, S. M., & Nichols, T. E. (2014). Permutation inference for the general linear model. NeuroImage, 92, 381–397.

Woo, C. W., Krishnan, A., & Wager, T. D. (2014). Cluster-extent based thresholding in fMRI analyses: pitfalls and recommendations. NeuroImage, 91, 412–419.

Wood, E. R., Dudchenko, P. A., Robitsek, R. J., & Eichenbaum, H. (2000). Hippocampal neurons encode information about different types of memory episodes occurring in the same location. Neuron, 27(3), 623–633.

Xu, Y., Vignali, L., Sigismondi, F., Crepaldi, D., Bottini, R., & Collignon, O. (2023). Similar object shape representation encoded in the inferolateral occipitotemporal cortex of sighted and early blind people. PLOS Biology, 21(7), e3001930.

Yeo, B. T., Krienen, F. M., Sepulcre, J., Sabuncu, M. R., Lashkari, D., Hollinshead, M., … & Buckner, R. L. (2011). The organization of the human cerebral cortex estimated by intrinsic functional connectivity. Journal of Neurophysiology, 106(3), 1125–1165.

Yoon, K., Buice, M. A., Barry, C., Hayman, R., Burgess, N., & Fiete, I. R. (2013). Specific evidence of low-dimensional continuous attractor dynamics in grid cells. Nature Neuroscience, 16(8), 1077–1084.

