## Supplementary material for "Neural Dissociation of Cognitive Maps and Graphs in the Human Brain": Figures S1-S7; Table S1

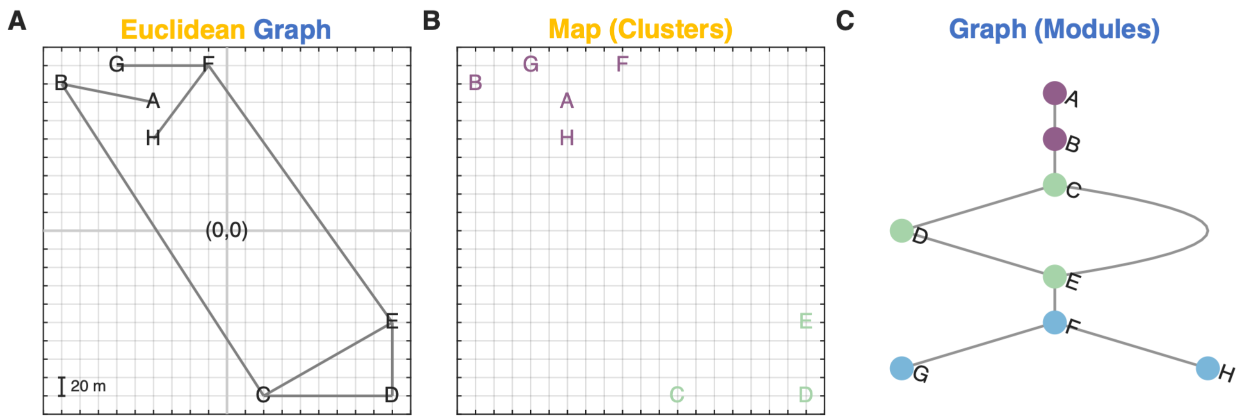


**Figure S1. Orthogonal design of the Euclidean graph for the virtual environment.**

**(A)** Euclidean graph serving as the blueprint for the virtual environment. Eight locations (A–H) located in a two-dimensional Euclidean space are interconnected. The origin of the coordinate system was defined at the centre of the 400 m × 400 m environment, and the basic spatial unit is 20 m.

**(B)** Clusters of the Euclidean map. K-means clustering of the Euclidean coordinates identified two clusters that can be separated by boundaries aligned with the lower-left–to–upper-right axis of the environment. These two clusters are denoted by different colours.

**(C)** Modules of the graph structure, displayed using a layered layout. Modularity-based community detection revealed three modules that can be separated by boundaries aligned with the upper-left–to–lower-right axis of the environment. These three modules are denoted by different colours.

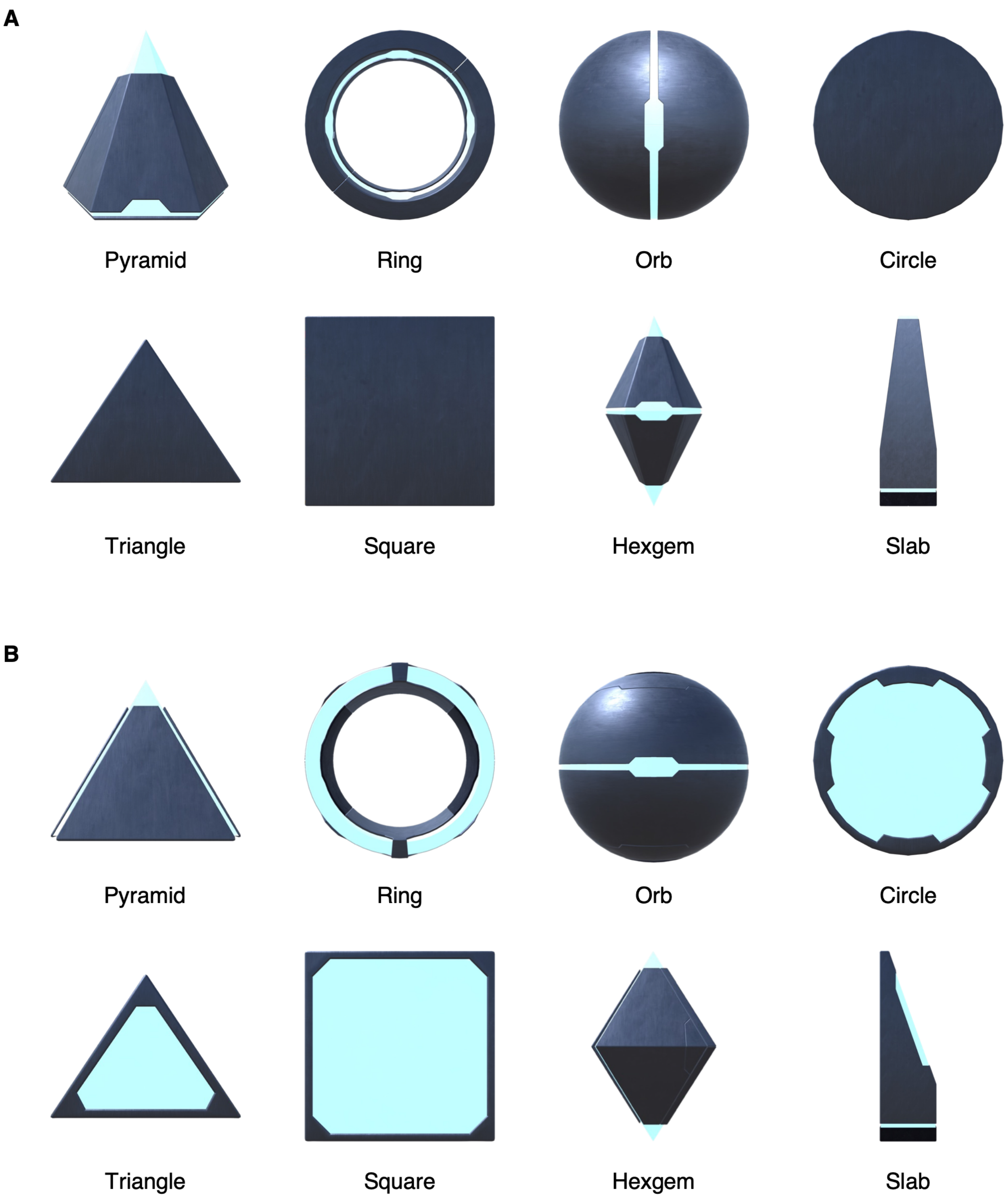


**Figure S2. Representative perspective views of sci-fi–style monuments.**

**(A)** Representative perspective views of the eight monuments marking the eight locations in the virtual environment. These monuments served as cue stimuli for participants to recall their positions within the cognitive map and graph during fMRI scanning.

**(B)** Representative perspective views of the monuments selected as visual analogues of those shown in panel (A). These monuments served as oddball stimuli in the fMRI experiment.

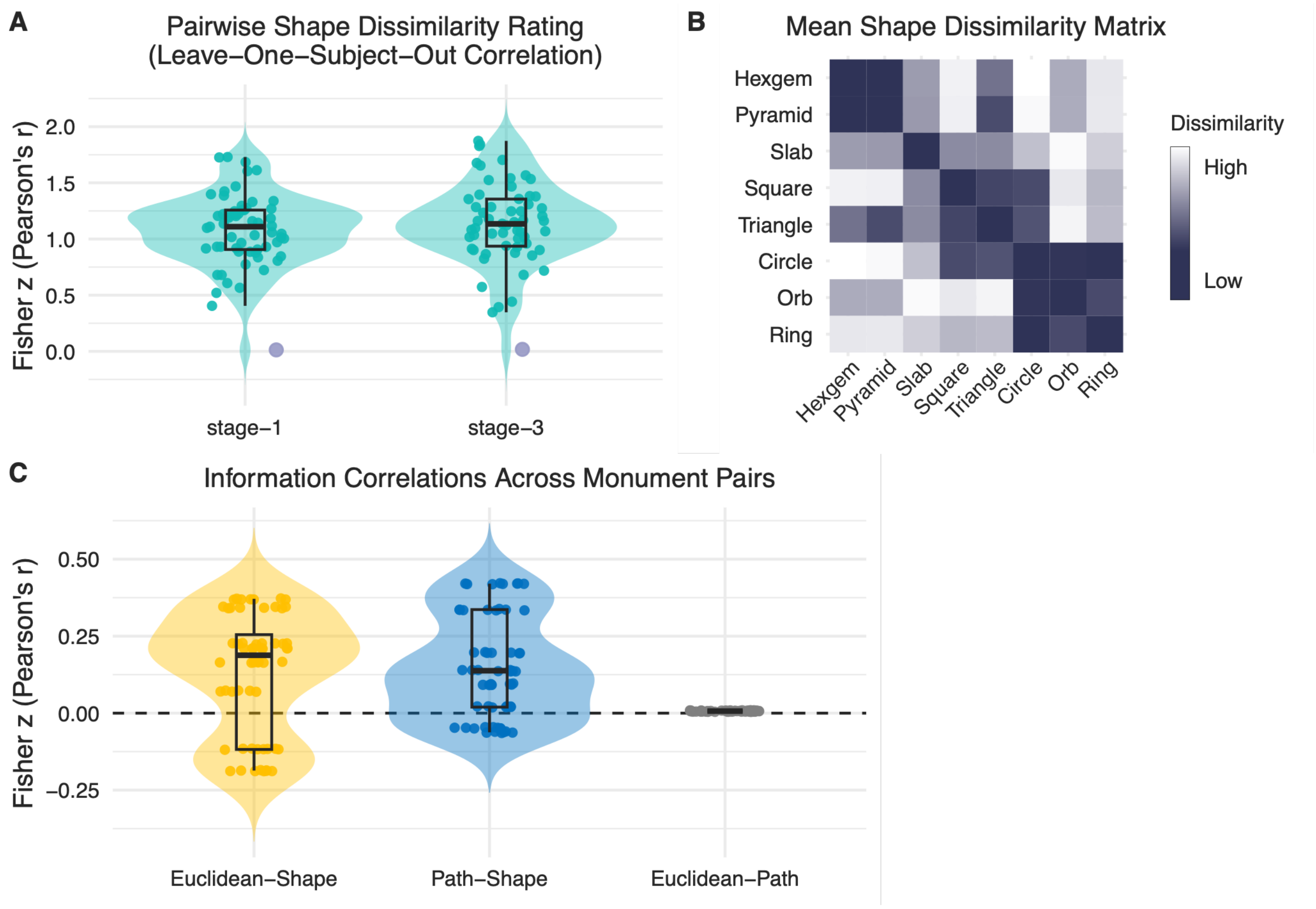


**Figure S3. Pairwise shape dissimilarity rating of the monuments.**

**(A)** To assess the consistency of pairwise shape ratings across participants, we computed leave-one-participant-out Pearson correlations between each participant's shape ratings and the mean shape ratings of all remaining participants. The resulting correlation coefficients were Fisher *z*-transformed before outlier detection. The same participant was identified as an outlier at both Stages 1 and 3, with a Fisher *z*-transformed correlation below the first quartile by more than 1.5 times the interquartile range (purple dot).

**(B)** The participant-averaged shape dissimilarity matrix from Stage 3, computed after removal of the outlier participant, was used as the shape dissimilarity matrix for the representational similarity analysis. The matrix exhibited a clear clustering structure, with stimuli largely grouped into three categories: pyramidal, polygonal, and rounded shapes.

**(C)** Pearson correlations among the Euclidean distance, shortest-path distance, and shape dissimilarity matrices across location pairs for each participant. Although the assignment of the eight monuments to the eight locations was counterbalanced across participants using a Latin square design to minimize potential influence of monument shape on representations of cognitive maps and graphs, the shape dissimilarity matrix nevertheless showed weak positive correlations with both Euclidean and shortest-path distances on average.

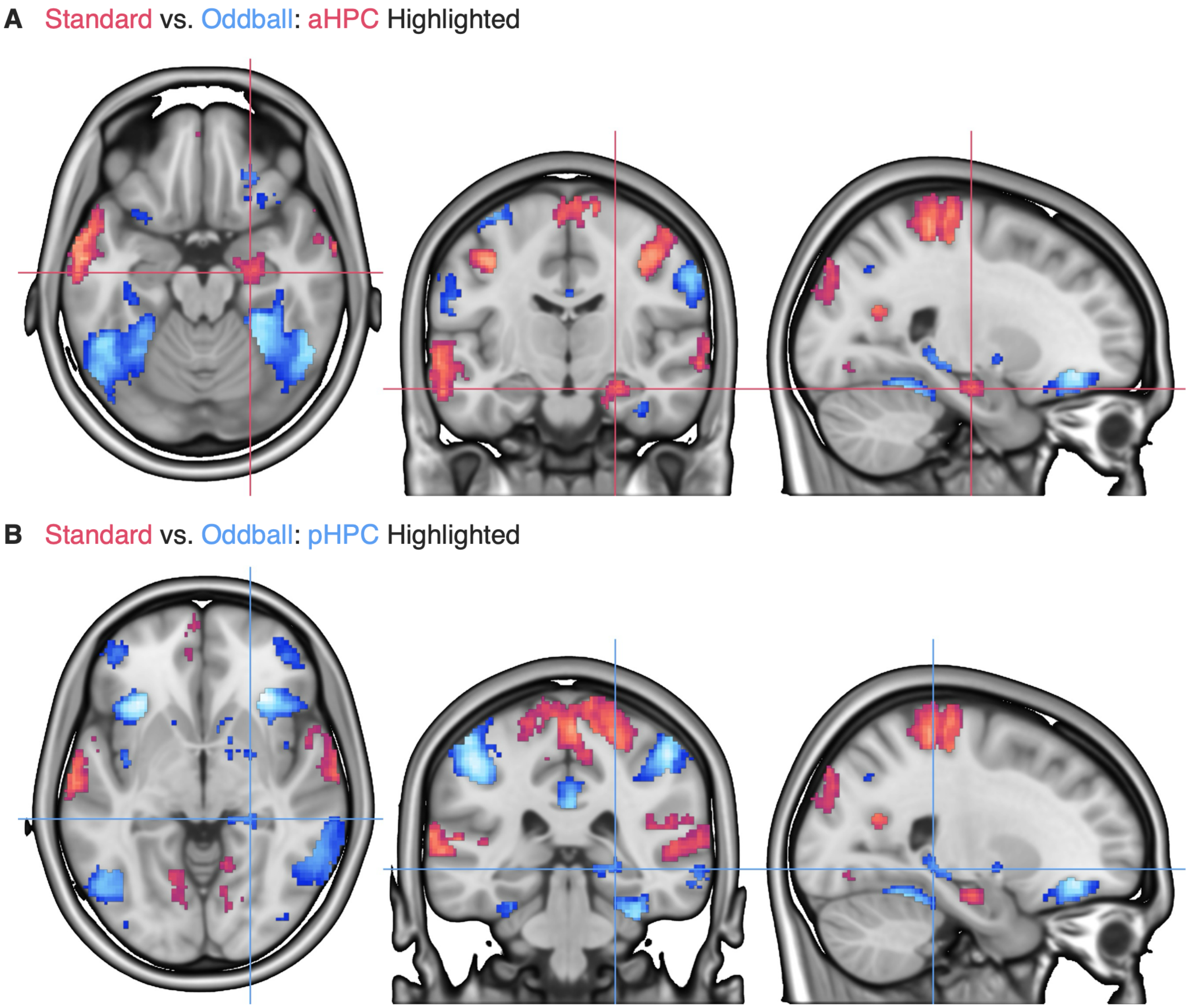


**Figure S4. Task-activation contrasts between the standard and oddball conditions, with hippocampal effects highlighted.** The z-statistic maps were thresholded using a voxel-wise cluster-forming threshold of $p < 0.001$ and a whole-brain cluster-level FWE-corrected threshold of $p < 0.05$.

**(A)** The anterior hippocampus (aHPC) is highlighted. The crosshair indicates MNI coordinates $x = 24$, $y = -15$, $z = -19$.

**(B)** The posterior hippocampus (pHPC) is highlighted. The crosshair indicates MNI coordinates $x = 24$, $y = -34$, $z = -5$.

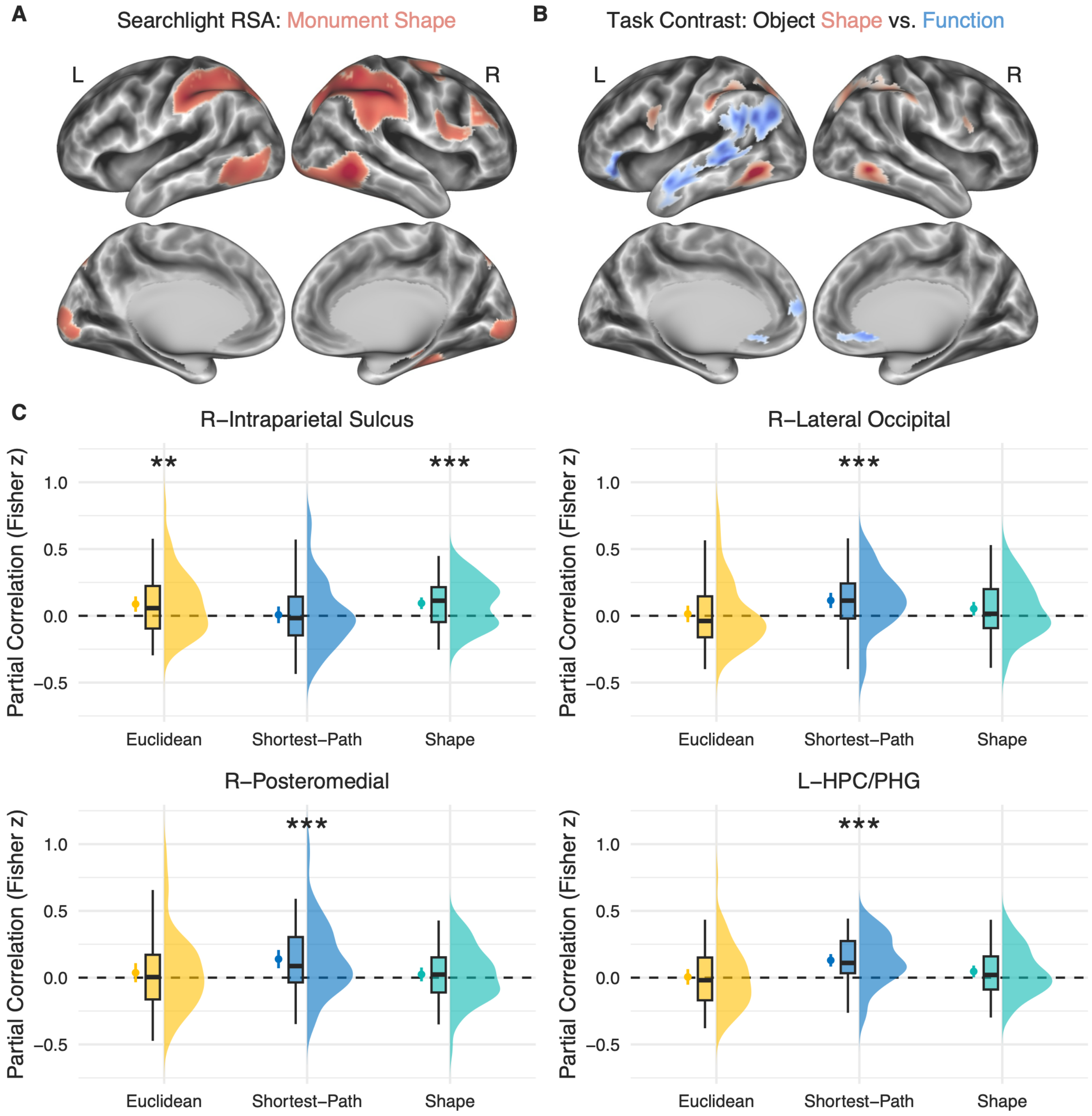


**Figure S5. Monument shape representations and unique contributions of Euclidean distance, shortest-path distance, and shape dissimilarity.**

**(A)** Whole-brain searchlight results associated with monument shape, based on the shape-dissimilarity matrix shown in Figure S3B. The *z*-statistic maps were thresholded using a voxel-wise cluster-forming threshold of $p < 0.001$ and a whole-brain cluster-level FWE-corrected threshold of $p < 0.05$. The spatial distribution of the effects broadly resembles the activation pattern observed in the independent shape-processing study shown in panel B.

**(B)** Task-related fMRI activation for Object Shape > Object Function from an independent study. Reproduced from Xu et al. (2023) under the Creative Commons Attribution (CC BY) License. Participants ($N = 48$) judged either the shape of an object (e.g., *Is the plate round?*) or its function (e.g., *Is the plate used for eating?*). The $z$-statistic maps were thresholded using a voxel-wise cluster-forming threshold of $p < 0.001$ and a whole-brain cluster-level FWE-corrected threshold of $p < 0.05$.

**(C)** Post-hoc robustness analysis within the ROIs identified in the primary whole-brain searchlight analysis. Partial correlations were computed for Euclidean distance, shortest-path distance, and shape dissimilarity while controlling for the other two models, to assess whether the originally observed representational effects remained after accounting for shared variance among the model RDMs. Points indicate group means across participants, and error bars indicate 95% confidence intervals. $p$ values were Holm-adjusted across the three information models within each ROI. ** $p_{\text{Holm}} < 0.01$; *** $p_{\text{Holm}} < 0.001$.

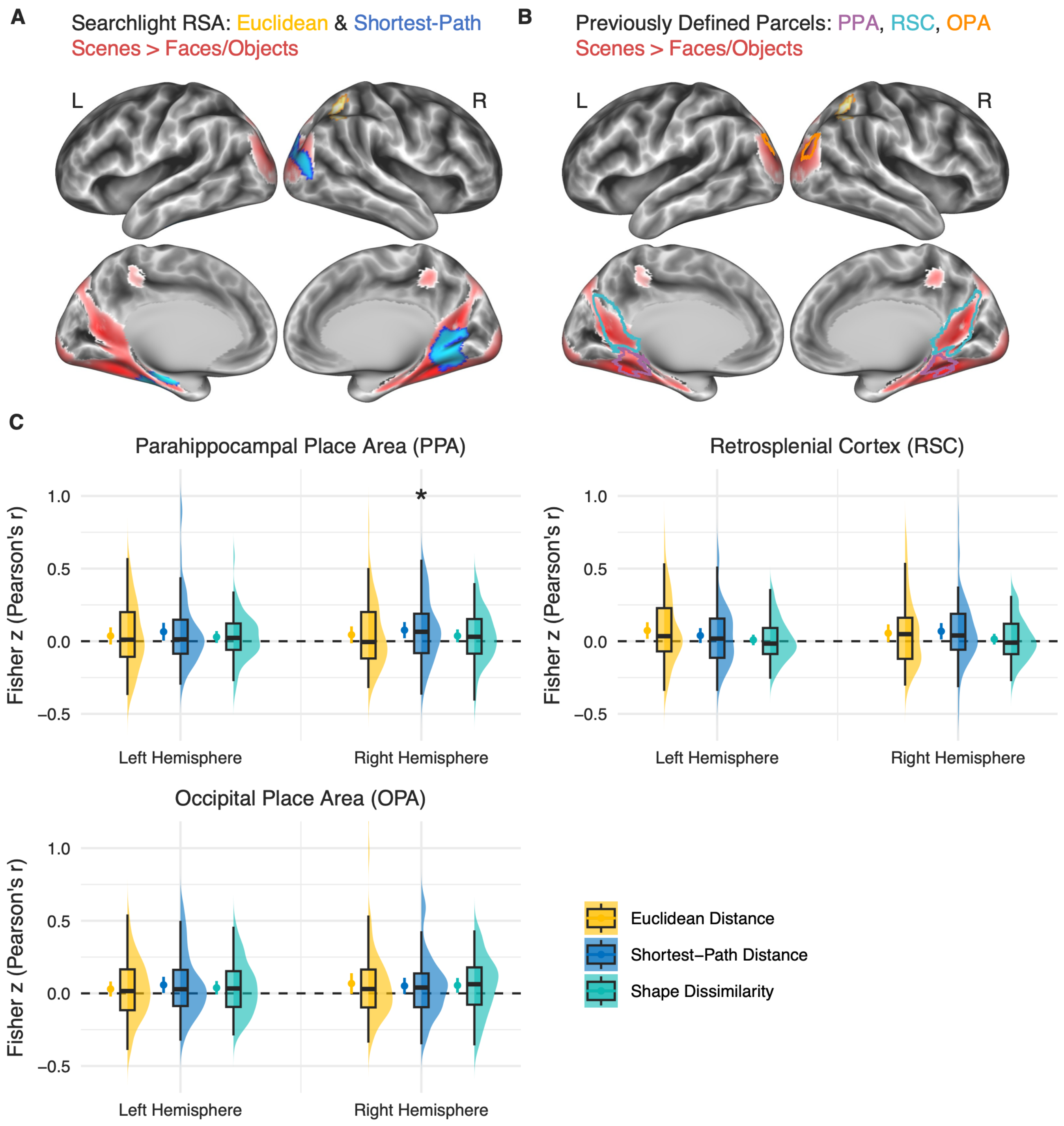


**Figure S6. Scene-selective regions and their representational effects.**

**(A)** Overlap between the whole-brain searchlight effects of Euclidean distance and shortest-path distance and scene-selective regions functionally defined using the contrast scenes > faces/objects. Both the searchlight and functional-localizer $z$-statistic maps were thresholded using a voxel-wise cluster-forming threshold of $p < 0.001$ and a whole-brain cluster-level FWE-corrected threshold of $p < 0.05$.

**(B)** Overlap between previously defined scene-selective parcels (Julian et al., 2012) and the scene-selective regions identified using the current functional localizer (scenes > faces/objects). The previously defined parcels showed substantial spatial overlap with the functionally localized regions. PPA = parahippocampal place area; RSC = retrosplenial cortex; OPA = occipital place area.

**(C)** Representational effects within each scene-selective region, shown separately for the left and right hemispheres. Euclidean distance, shortest-path distance, and shape dissimilarity were tested against zero within each hemisphere. Points indicate group means across participants, and error bars indicate 95% confidence intervals. $p$ values were Holm-adjusted across the six comparisons (three information models × two hemispheres) within each ROI. * $p_{\mathrm{Holm}} < 0.05$.

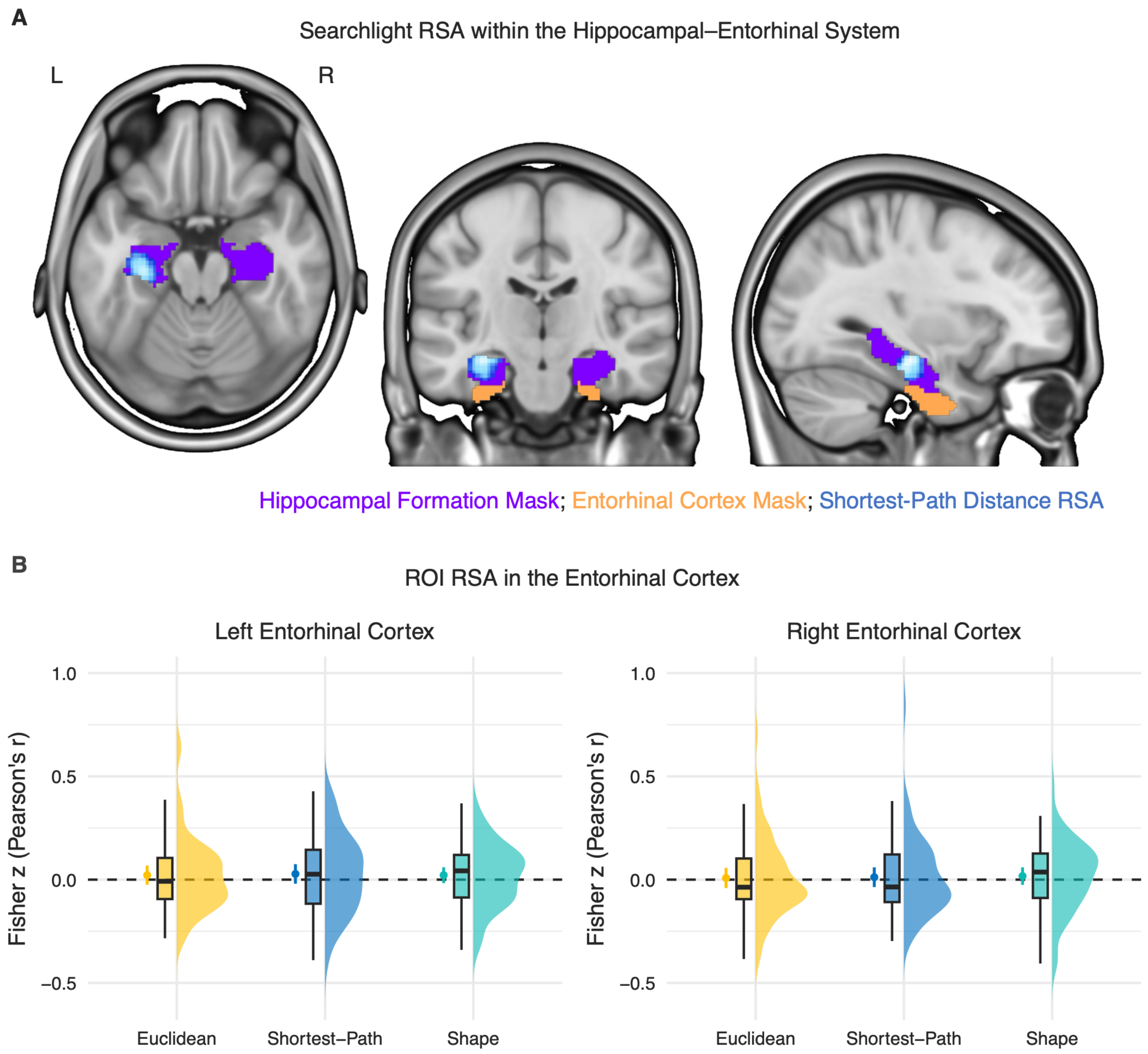


**Figure S7. Representational similarity analysis within the hippocampal–entorhinal system.**

**(A)** Searchlight RSA within the hippocampal–entorhinal system. Euclidean distance, shortest-path distance, and shape dissimilarity were tested. A significant effect was observed only for shortest-path distance, confined to the hippocampal formation (purple), with no significant effect in the entorhinal cortex (orange). Searchlight results are shown as *z*-statistic maps thresholded at a voxel-wise cluster-forming threshold of $p < 0.001$ and a cluster-level FWE-corrected threshold of $p < 0.05$.

**(B)** ROI RSA in the entorhinal cortex. Neither the left nor right entorhinal cortex showed significant representations of Euclidean distance, shortest-path distance, or shape dissimilarity. Points indicate group means across participants, and error bars indicate 95% confidence intervals. All $p$s > 0.25.

**Table S1. Virtual environment learning**

| Days | Stages | Tasks | Post-Learning Tests[1] |
|---|---|---|---|
| Day 1 | Stage 1: Familiarization | Free Exploration<br>(5 minutes) | Pairwise Rating of Euclidean Distance<br>Pairwise Rating of Shortest-Path Distance<br>Pairwise Rating of Shape Dissimilarity |
| | Stage 2: Main Learning | Goal-Directed Navigation<br>(~ 1 hour) | No post-learning tests |
| | Stage 3: Wrap-Up | Free Exploration<br>(no time limit) | Pairwise Rating of Euclidean Distance<br>Spatial Arrangement Task<br>Pairwise Rating of Shortest-Path Distance<br>Shortest-Path Selection<br>Pairwise Rating of Shape Dissimilarity |
| Day 2 | Stage 4: Warm-Up | Free Exploration<br>(no time limit) | No post-learning tests |

[1] Tests assessing knowledge of the cognitive map are shown in yellow, those assessing knowledge of the cognitive graph are shown in blue, and those assessing knowledge of the monument shape are shown in turquoise.

### Supplemental methods

Results included in this manuscript come from preprocessing performed using *fMRIPrep* 22.0.1 (Esteban, Markiewicz, et al. (2018); Esteban, Blair, et al. (2018); RRID:SCR_016216), which is based on *Nipype* 1.8.4 (K. Gorgolewski et al. (2011); K. J. Gorgolewski et al. (2018); RRID:SCR_002502).

*Preprocessing of $B_0$ inhomogeneity mappings*

A total of 3 fieldmaps were found available within the input BIDS structure for this particular subject. A $B_0$-nonuniformity map (or *fieldmap*) was estimated based on two (or more) echo-planar imaging (EPI) references with topup (Andersson, Skare, and Ashburner (2003); FSL 6.0.5.1:57b01774).

*Anatomical data preprocessing*

A total of 1 T1-weighted (T1w) images were found within the input BIDS dataset.The T1-weighted (T1w) image was corrected for intensity non-uniformity (INU) with N4BiasFieldCorrection (Tustison et al. 2010), distributed with ANTs 2.3.3 (Avants et al. 2008, RRID:SCR_004757), and used as T1w-reference throughout the workflow. The T1w-reference was then skull-stripped with a *Nipype* implementation of the antsBrainExtraction.sh workflow (from ANTs), using OASIS30ANTs as target template. Brain tissue segmentation of cerebrospinal fluid (CSF), white-matter (WM) and gray-matter (GM) was performed on the brain-extracted T1w using fast (FSL 6.0.5.1:57b01774, RRID:SCR_002823, Zhang, Brady, and Smith 2001). Brain surfaces were reconstructed using recon-all (FreeSurfer 7.2.0, RRID:SCR_001847, Dale, Fischl, and Sereno 1999), and the brain mask estimated previously was refined with a custom variation of the method to reconcile ANTs-derived and FreeSurfer-derived segmentations of the cortical gray-matter of Mindboggle (RRID:SCR_002438, Klein et al. 2017). Volume-based spatial normalization to two standard spaces (MNI152NLin6Asym, MNI152NLin2009cAsym) was performed through nonlinear registration with antsRegistration (ANTs 2.3.3), using brain-extracted versions of both T1w reference and the T1w template. The following templates were selected for spatial normalization: *FSL's MNI ICBM 152 non-linear 6th Generation Asymmetric Average Brain Stereotaxic Registration Model* [Evans et al. (2012), RRID:SCR_002823; TemplateFlow ID: MNI152NLin6Asym], *ICBM 152 Nonlinear Asymmetrical template version 2009c* [Fonov et al. (2009), RRID:SCR_008796; TemplateFlow ID: MNI152NLin2009cAsym].

*Functional data preprocessing*

For each of the 10 BOLD runs found per subject (across all tasks and sessions), the following preprocessing was performed. First, a reference volume and its skull-stripped version were generated using a custom methodology of *fMRIPrep*. Head-motion parameters with respect to the BOLD reference (transformation matrices, and six corresponding rotation and translation parameters) are estimated before any spatiotemporal filtering using mcflirt (FSL 6.0.5.1:57b01774, Jenkinson et al. 2002). The estimated *fieldmap* was then aligned with rigid-registration to the target EPI (echo-planar imaging) reference run. The field coefficients were mapped on to the reference EPI using the transform. BOLD runs were slice-time corrected to 0.451s (0.5 of slice acquisition range 0s-0.902s) using 3dTshift from AFNI (Cox and Hyde 1997, RRID:SCR_005927). The BOLD reference was then co-registered to the T1w reference using bbregister (FreeSurfer) which implements boundary-based registration (Greve and Fischl 2009). Co-registration was configured with six degrees of freedom. Several confounding time-series were calculated based on the *preprocessed BOLD*: framewise displacement (FD), DVARS and three region-wise global signals. FD was computed using two formulations following Power (absolute sum of relative motions, Power et al. (2014)) and Jenkinson (relative root mean square displacement between affines, Jenkinson et al. (2002)). FD and DVARS are calculated for each functional run, both using their implementations in *Nipype* (following the definitions by Power et al. 2014). The three global signals are extracted within the CSF, the WM, and the whole-brain masks. Additionally, a set of physiological regressors were extracted to allow for component-based noise correction(*CompCor*, Behzadi et al. 2007). Principal components are estimated after high-pass filtering the *preprocessed BOLD* time-series (using a discrete cosine filter with 128s cut-off) for the two *CompCor* variants: temporal (tCompCor) and anatomical (aCompCor). tCompCor components are then calculated from the top 2% variable voxels within the brain mask. For aCompCor, three probabilistic masks (CSF, WM and combined CSF+WM) are generated in anatomical space. The implementation differs from that of Behzadi et al. in that instead of eroding the masks by 2 pixels on BOLD space, a mask of pixels that likely contain a volume fraction of GM is subtracted from the aCompCor masks. This mask is obtained by dilating a GM mask extracted from the FreeSurfer's *aseg* segmentation, and it ensures components are not extracted from voxels containing a minimal fraction of GM. Finally, these masks are resampled into BOLD space and binarized by thresholding at 0.99 (as in the original implementation). Components are also calculated separately within the WM and CSF masks. For each CompCor decomposition, the *k* components with the largest singular values are retained, such that the retained components' time series are sufficient to explain 50 percent of variance across the nuisance mask (CSF, WM, combined, or temporal). The remaining components are dropped from consideration. The head-motion estimates calculated in the correction step were also placed within the corresponding confounds file. The confound time series derived from head motion estimates and global signals were expanded with the inclusion of temporal derivatives and quadratic terms for each (Satterthwaite et al. 2013). Frames that exceeded a threshold of 0.5 mm FD or 1.5 standardized DVARS were annotated as motion outliers. Additional nuisance timeseries are calculated by means of principal components analysis of the signal found within a thin band (*crown*) of voxels around the edge of the brain, as proposed by (Patriat, Reynolds, and Birn 2017). The BOLD time-series were resampled into standard space, generating a *preprocessed BOLD run in MNI152NLin6Asym space*. First, a reference volume and its skull-stripped version were generated using a custom methodology of *fMRIPrep*. The BOLD time-series were resampled onto the following surfaces (FreeSurfer reconstruction nomenclature): *fsaverage5*, *fsaverage6*, *fsaverage*. Automatic removal of motion artifacts using independent component analysis (ICA-AROMA, Pruim et al. 2015) was performed on the *preprocessed BOLD on MNI space* time-series after removal of non-steady state volumes and spatial smoothing with an isotropic, Gaussian kernel of 6mm FWHM (full-width half-maximum). Corresponding "non-aggresively" denoised runs were produced after such smoothing. Additionally, the "aggressive" noise-regressors were collected and placed in the corresponding confounds file. *Grayordinates* files (Glasser et al. 2013) containing 91k samples were also generated using the highest-resolution fsaverage as intermediate standardized surface space. All resamplings can be performed with *a single interpolation step* by composing all the pertinent transformations (i.e. head-motion transform matrices, susceptibility distortion correction when available, and co-registrations to anatomical and output spaces). Gridded (volumetric) resamplings were performed usingantsApplyTransforms (ANTs), configured with Lanczos interpolation to minimize the smoothing effects of other kernels (Lanczos 1964). Non-gridded (surface) resamplings were performed using mri_vol2surf (FreeSurfer).

Many internal operations of *fMRIPrep* use *Nilearn* 0.9.1 (Abraham et al. 2014, RRID:SCR_001362), mostly within the functional processing workflow. For more details of the pipeline, see the section corresponding to workflows in *fMRIPrep*'s documentation.

*Copyright Waiver*

The above boilerplate text was automatically generated by fMRIPrep with the express intention that users should copy and paste this text into their manuscripts *unchanged*. It is released under the CC0 license.

*References*


Abraham, Alexandre, Fabian Pedregosa, Michael Eickenberg, Philippe Gervais, Andreas Mueller, Jean Kossaifi, Alexandre Gramfort, Bertrand Thirion, and Gael Varoquaux. 2014. "Machine Learning for Neuroimaging with Scikit-Learn." *Frontiers in Neuroinformatics* 8. https://doi.org/10.3389/fninf.2014.00014.

Andersson, Jesper L. R., Stefan Skare, and John Ashburner. 2003. "How to Correct Susceptibility Distortions in Spin-Echo Echo-Planar Images: Application to Diffusion Tensor Imaging." *NeuroImage* 20 (2): 870–88. https://doi.org/10.1016/S1053-8119(03)00336-7.

Avants, B. B., C. L. Epstein, M. Grossman, and J. C. Gee. 2008. "Symmetric Diffeomorphic Image Registration with Cross-Correlation: Evaluating Automated Labeling of Elderly and Neurodegenerative Brain." *Medical Image Analysis* 12 (1): 26–41. https://doi.org/10.1016/j.media.2007.06.004.

Behzadi, Yashar, Khaled Restom, Joy Liau, and Thomas T. Liu. 2007. "A Component Based Noise Correction Method (CompCor) for BOLD and Perfusion Based fMRI."*NeuroImage* 37 (1): 90–101. https://doi.org/10.1016/j.neuroimage.2007.04.042.

Cox, Robert W., and James S. Hyde. 1997. "Software Tools for Analysis and Visualization of fMRI Data." *NMR in Biomedicine* 10 (4-5): 171–78. https://doi.org/10.1002/(SICI)1099-1492(199706/08)10:4/5<171::AID-NBM453>3.0.CO;2-L.

Dale, Anders M., Bruce Fischl, and Martin I. Sereno. 1999. "Cortical Surface-Based Analysis: I. Segmentation and Surface Reconstruction." *NeuroImage* 9 (2): 179–94. https://doi.org/10.1006/nimg.1998.0395.

Esteban, Oscar, Ross Blair, Christopher J. Markiewicz, Shoshana L. Berleant, Craig Moodie, Feilong Ma, Ayse Ilkay Isik, et al. 2018. "fMRIPrep 22.0.1." *Software*. https://doi.org/10.5281/zenodo.852659.

Esteban, Oscar, Christopher Markiewicz, Ross W Blair, Craig Moodie, Ayse Ilkay Isik, Asier Erramuzpe Aliaga, James Kent, et al. 2018. "fMRIPrep: A Robust Preprocessing Pipeline for Functional MRI." *Nature Methods*. https://doi.org/10.1038/s41592-018-0235-4.

Evans, AC, AL Janke, DL Collins, and S Baillet. 2012. "Brain Templates and Atlases." *NeuroImage* 62 (2): 911–22. https://doi.org/10.1016/j.neuroimage.2012.01.024.

Fonov, VS, AC Evans, RC McKinstry, CR Almli, and DL Collins. 2009. "Unbiased Nonlinear Average Age-Appropriate Brain Templates from Birth to Adulthood."*NeuroImage* 47, Supplement 1: S102. https://doi.org/10.1016/S1053-8119(09)70884-5.

Glasser, Matthew F., Stamatios N. Sotiropoulos, J. Anthony Wilson, Timothy S. Coalson, Bruce Fischl, Jesper L. Andersson, Junqian Xu, et al. 2013. "The Minimal Preprocessing Pipelines for the Human Connectome Project." *NeuroImage*, Mapping the connectome, 80: 105–24. https://doi.org/10.1016/j.neuroimage.2013.04.127.

Gorgolewski, K., C. D. Burns, C. Madison, D. Clark, Y. O. Halchenko, M. L. Waskom, and S. Ghosh. 2011. "Nipype: A Flexible, Lightweight and Extensible Neuroimaging Data Processing Framework in Python." *Frontiers in Neuroinformatics* 5: 13. https://doi.org/10.3389/fninf.2011.00013.

Gorgolewski, Krzysztof J., Oscar Esteban, Christopher J. Markiewicz, Erik Ziegler, David Gage Ellis, Michael Philipp Notter, Dorota Jarecka, et al. 2018. “Nipype.”*Software*. https://doi.org/10.5281/zenodo.596855.

Greve, Douglas N, and Bruce Fischl. 2009. “Accurate and Robust Brain Image Alignment Using Boundary-Based Registration.” *NeuroImage* 48 (1): 63–72. https://doi.org/10.1016/j.neuroimage.2009.06.060.

Jenkinson, Mark, Peter Bannister, Michael Brady, and Stephen Smith. 2002. “Improved Optimization for the Robust and Accurate Linear Registration and Motion Correction of Brain Images.” *NeuroImage* 17 (2): 825–41. https://doi.org/10.1006/nimg.2002.1132.

Klein, Arno, Satrajit S. Ghosh, Forrest S. Bao, Joachim Giard, Yrjö Häme, Eliezer Stavsky, Noah Lee, et al. 2017. “Mindboggling Morphometry of Human Brains.” *PLOS Computational Biology* 13 (2): e1005350. https://doi.org/10.1371/journal.pcbi.1005350.

Lanczos, C. 1964. “Evaluation of Noisy Data.” *Journal of the Society for Industrial and Applied Mathematics Series B Numerical Analysis* 1 (1): 76–85. https://doi.org/10.1137/0701007.

Patriat, Rémi, Richard C. Reynolds, and Rasmus M. Birn. 2017. “An Improved Model of Motion-Related Signal Changes in fMRI.” *NeuroImage* 144, Part A (January): 74–82. https://doi.org/10.1016/j.neuroimage.2016.08.051.

Power, Jonathan D., Anish Mitra, Timothy O. Laumann, Abraham Z. Snyder, Bradley L. Schlaggar, and Steven E. Petersen. 2014. “Methods to Detect, Characterize, and Remove Motion Artifact in Resting State fMRI.” *NeuroImage* 84 (Supplement C): 320–41. https://doi.org/10.1016/j.neuroimage.2013.08.048.

Pruim, Raimon H. R., Maarten Mennes, Daan van Rooij, Alberto Llera, Jan K. Buitelaar, and Christian F. Beckmann. 2015. “ICA-AROMA: A Robust ICA-Based Strategy for Removing Motion

Artifacts from fMRI Data." *NeuroImage* 112 (Supplement C): 267–77. https://doi.org/10.1016/j.neuroimage.2015.02.064.

Satterthwaite, Theodore D., Mark A. Elliott, Raphael T. Gerraty, Kosha Ruparel, James Loughead, Monica E. Calkins, Simon B. Eickhoff, et al. 2013. "An improved framework for confound regression and filtering for control of motion artifact in the preprocessing of resting-state functional connectivity data." *NeuroImage* 64 (1): 240–56. https://doi.org/10.1016/j.neuroimage.2012.08.052.

Tustison, N. J., B. B. Avants, P. A. Cook, Y. Zheng, A. Egan, P. A. Yushkevich, and J. C. Gee. 2010. "N4itk: Improved N3 Bias Correction." *IEEE Transactions on Medical Imaging* 29 (6): 1310–20. https://doi.org/10.1109/TMI.2010.2046908.

Zhang, Y., M. Brady, and S. Smith. 2001. "Segmentation of Brain MR Images Through a Hidden Markov Random Field Model and the Expectation-Maximization Algorithm." *IEEE Transactions on Medical Imaging* 20 (1): 45–57. https://doi.org/10.1109/42.906424.
